# Geometry-enhanced protein language modeling enables discovery of novel antibiotic resistance genes

**DOI:** 10.64898/2026.04.05.716581

**Authors:** Xingqiao Lin, Jiahui Guan, Yue Hong, Yilian Guo, Yutao Yang, Peilin Xie, Zhihao Zhao, Xingchen Liu, Yixian Huang, Yujing Ye, Yun Tang, Tzong-Yi Lee, Ying-Chih Chiang, Leyi Wei, Xiangrong Liu, Junwen Wang, Yi Pan, Jijun Tang, Yao Pei, Lantian Yao

## Abstract

The global antibiotic resistome remains largely unexplored, not because antibiotic resistance genes (ARGs) are rare in the environment, but because many are evolutionarily distant from known ARGs. Current computational approaches primarily rely on sequence homology, and thus miss distant homologues. We develop GeoARG, a geometry-enhanced framework that integrates structural features with protein language models through knowledge distillation, enabling efficient large-scale screening using sequence input alone. Across multiple benchmarks, GeoARG substantially improves the detection of remotely homologous ARGs, particularly under low sequence identity and fragmented conditions. Large-scale metagenomic analysis uncovers 1,485 high-confidence ARG candidates that are highly divergent from known ARGs, expanding the phylogenetic and functional landscape of the resistome. Structural analyses further show that these candidates preserve active-site geometry and maintain stable ligand-binding configurations consistent with known resistance mechanisms. These results demonstrate that geometric constraints enable systematic expansion of the resistome and facilitate the discovery of evolutionarily distant yet functionally conserved genes. A public web server is available at https://ycclab.cuhk.edu.cn/GeoARG/.

## 1 Introduction

Antimicrobial resistance (AMR) is a rapidly escalating global health crisis projected to cause up to 10 million deaths per year by 2050 and is currently responsible for approximately 700,000 deaths annually. and is projected to reach 10 million by 2050. [1–3]. At the molecular level, AMR is mediated by antibiotic resistance genes (ARGs), whose sequences diversify extensively through horizontal gene transfer, point mutations, and selective pressure from antibiotics [4–6]. Consequently, large-scale sequencing of environmental and clinical microbiomes routinely uncovers putative ARGs with little similarity to current reference databases [7, 8], suggesting that a substantial fraction of the functional resistome remains uncharacterized. Traditional ARG identification relies on sequence alignment against curated reference databases using tools such as BLAST [9] or DIAMOND [10]. For instance, Jia et al.[11] proposed the Comprehensive Antibiotic Resistance Database (CARD) and the Resistance Gene Identifier (RGI), which apply predefined similarity thresholds and DIAMOND [10] for ARG detection and annotation [12, 13]. While these approaches are effective for identifying closely related sequences, their performance is constrained by database coverage and fixed similarity cutoffs and therefore often fail to detect distantly related or novel ARGs. Moreover, choosing an appropriate similarity threshold across diverse ARG families is challenging and can further increase false-negative rates in large-scale metagenomic analyses [14, 15].

To overcome these limitations, deep learning approaches have been developed for ARG annotation, where, unlike alignment-based methods, these tools learn features directly from sequence data without restriction by a strict similarity cutoff, thereby enhancing detection sensitivity [14–16]. For example, DeepARG uses neural networks trained on DIAMOND-generated features to extend detection beyond best-hit criteria [14]. HMD-ARG applies a hierarchical multi-task convolutional neural network to identify ARGs and predict their antibiotic classes [16], while ARGNet adopts a two-stage framework that combines an unsupervised autoencoder for ARG identification with a convolutional neural network for resistance type classification [15]. Collectively, these methods reduce reliance on predefined similarity thresholds and improve the identification of novel ARGs [15].

Despite these advances, several critical challenges remain for ARG detection. First, most existing models rely primarily on sequence features without explicitly modeling catalytic residues. Since ARG function is often determined by active-site residues [17, 18], the absence of structural information makes it difficult to distinguish functionally related enzyme families. Second, the high sequence divergence observed in metagenomic ARGs continues to make sequence-similarity-based detection insufficient and limited to the discovery of novel resistance determinants. Third, current frameworks require separate models for different input types or external contig context. For instance, DeepARG [14] and ARGNet [15] maintain distinct models for long versus short sequences. DRAMMA [19] requires nucleotide contig context as input, restricting its applicability when only protein sequences are available.

To address these limitations, we developed GeoARG, a geometry-enhanced framework for ARG identification and annotation that integrates three-dimensional structural information with sequence-based protein language modeling via knowledge distillation. GeoARG adopts a unified architecture capable of handling both long and short sequences across nucleotide and amino acid inputs, improving the detection of remotely homologous ARGs and enabling systematic discovery of novel resistance candidates in large-scale metagenomic datasets.

## 2 Results

### 2.1 Overview of GeoARG

GeoARG is a deep learning-based framework for identifying ARGs that are difficult to recover using sequence similarity alone, particularly in low-identity, fragmentary, and previously unannotated sequence settings commonly encountered in metagenomic screening. To address this challenge, GeoARG jointly models evolutionary context from protein sequences and catalytic-site geometry from three-dimensional structures, thereby improving sensitivity to remote homologs while maintaining scalability for large-scale deployment (Fig. 1a-d).

**Fig. 1.**
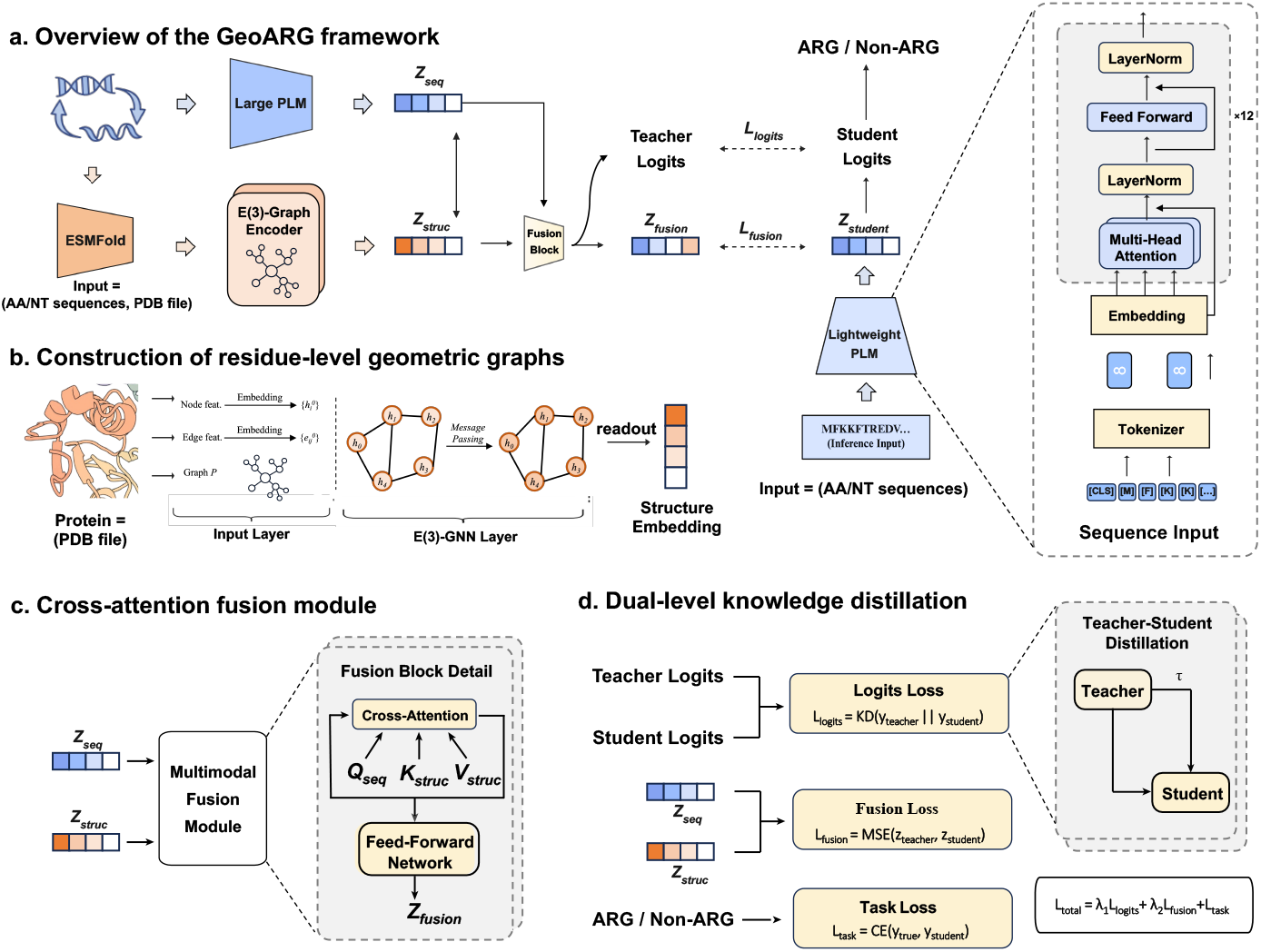
Architecture of GeoARG. **a** Overview of the GeoARG framework, in which sequence and structural representations are jointly learned in a multimodal teacher and distilled into a lightweight sequence-only student. **b** Construction of residue-level geometric graphs from predicted protein structures for structural encoding. **c** Crossattention fusion of sequence and structure representations in the teacher model. **d** Dual-level knowledge distillation from the multimodal teacher to the deployable student model.

The overall architecture of GeoARG consists of three coordinated modules. (1) A multimodal residue-level encoding module extracts complementary sequence and geometric representations from protein sequences and structures. (2) A fusion-and-distillation module integrates these teacher-level representations and transfers multimodal knowledge to a compact sequence-only student model. (3) A prediction module applies the distilled student representation to unified ARG classification tasks across different input forms. Together, these modules form an end-to-end framework for large-scale ARG identification and subtype annotation.

In the multimodal residue-level encoding module, amino acid sequences are first converted into contextual residue embeddings by a pretrained protein language model (PLM). In parallel, predicted protein structures are transformed into residue graphs, in which residues are treated as nodes and spatially proximal residue pairs are connected as edges. An E(3)-equivariant graph encoder then performs message passing on this graph to model residue interactions while preserving geometric relationships independent of global rotation and translation. Through this design, GeoARG captures local structural patterns and catalytic-site organization that are often only weakly reflected by sequence identity.

In the fusion-and-distillation module, the sequence and structure representations learned by the teacher are integrated through cross-attention to generate multimodal embeddings for ARG prediction. Knowledge is subsequently distilled from the teacher to the student at two levels. At the output level, the student is aligned with teacher logits to preserve predictive decision boundaries. At the representation level, the student is aligned with fused teacher embeddings to retain the internal geometry of the multimodal feature space. In this way, structural information informs training, whereas inference is ultimately performed using only a lightweight sequence-based student model, substantially reducing computational cost for large-scale screening.

In the prediction module, distilled student representations are aggregated and passed to task-specific classification heads for ARG prediction. The same student pipeline is applied across four practical input scenarios, namely long nucleotide sequences (LSnt), short nucleotide sequences (SSnt), long amino acid sequences (LSaa), and short amino acid sequences (SSaa), thereby avoiding separate model designs for different sequence types and lengths. Under this unified formulation, GeoARG supports both binary ARG identification and 36-class subtype annotation, improving deployment consistency while preserving sensitivity to short fragments and evolutionarily remote ARGs.

Following this design, we evaluated GeoARG at three levels: benchmark-level binary ARG identification, resistance subtype prediction, and downstream validation in independent metagenomic datasets. These analyses were intended to assess not only predictive accuracy, but also specificity under challenging negative backgrounds, sensitivity to divergent ARG families, and the practical utility of GeoARG for large-scale discovery. Detailed definitions and implementation of each module are provided in the “Methods”.

### 2.2 GeoARG-DB construction and benchmark design

Because performance in ARG prediction can be strongly influenced by the composition of negative backgrounds, we first established GeoARG-DB together with two complementary benchmark settings (Fig. 2). GeoARG-DB was assembled from seven public ARG resources. After cross-database harmonization, exclusion of SNP-mediated resistance records, and exact-sequence deduplication, the final collection contained 40,624 non-redundant proteins spanning 36 resistance classes (Fig. 2a,d). This curation step was intended to maximize resistance-class coverage while reducing annotation inconsistency across databases.

**Fig. 2.**
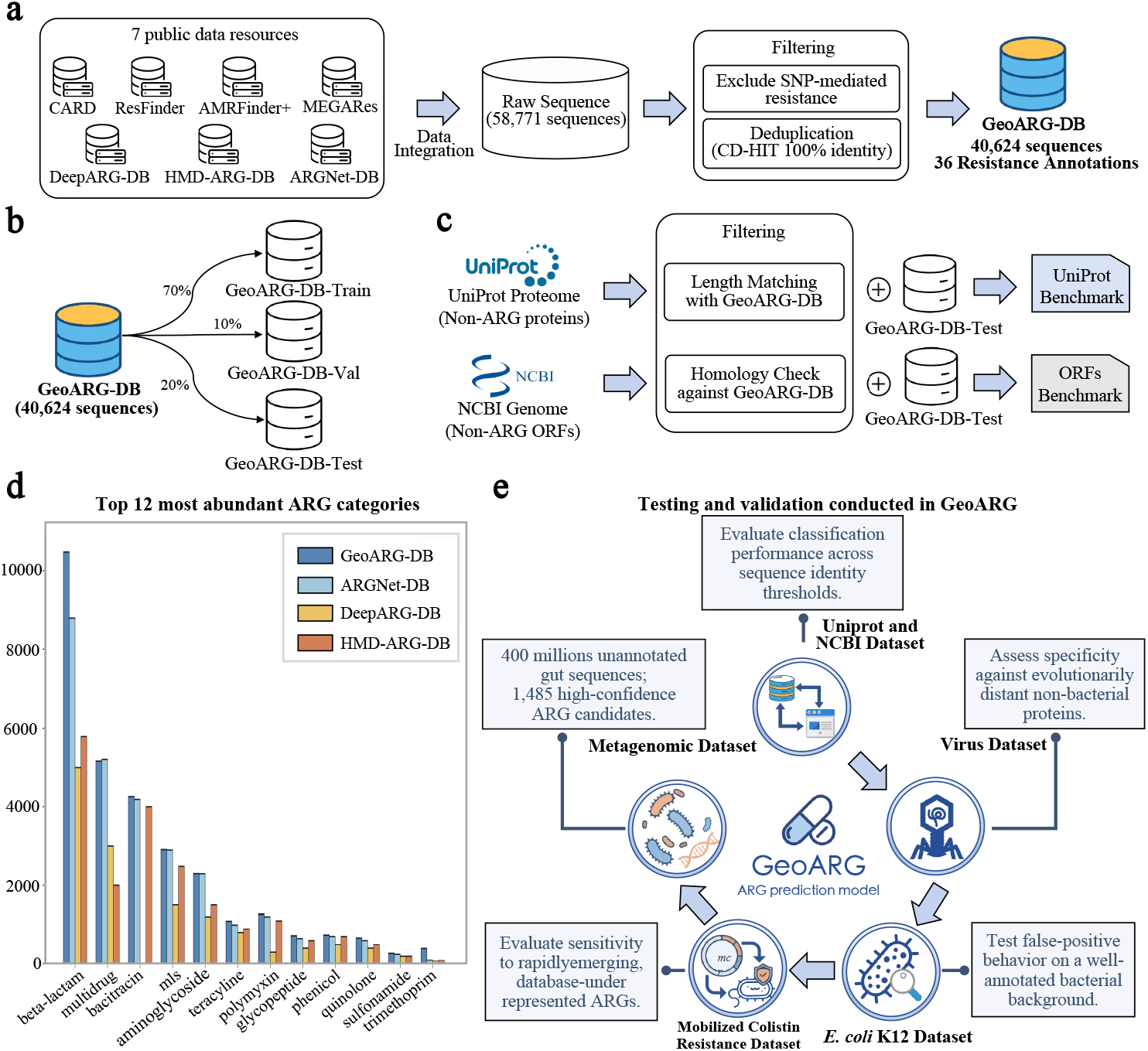
GeoARG-DB construction and benchmark design. **a** Integration and filtering of seven public ARG resources to construct GeoARG-DB, yielding 40,624 non-redundant sequences with 36 resistance annotations. **b** Partition of GeoARG-DB into training, validation, and test subsets (70/10/20%). **c** Construction of two primary benchmark settings by pairing the same positive test set with UniProt-derived or genome-derived ORF negatives after homology filtering. **d** Distribution of the most abundant resistance categories across representative ARG databases. **e** Independent validation datasets used to assess specificity, robustness to sequence divergence, and metagenomic discovery utility.

For model development, GeoARG-DB was divided into training (28,437, 70%), validation (4,062, 10%), and test (8,125, 20%) subsets (Fig. 2b; Supplementary Information Section S1.3). To evaluate robustness under different deployment scenarios, we fixed GeoARG-DB-Test as the positive set and generated two homology-filtered negative backgrounds: a UniProt bacterial protein background and a genome-derived ORF background from 1,000 NCBI bacterial genomes [20] (Fig. 2c).

These two settings were designed to capture complementary application scenarios. The UniProt benchmark approximates proteome-scale screening against curated bacterial proteins, whereas the ORF benchmark mimics gene-level identification in genomic or metagenomic assemblies, where fragmented and less curated inputs are more common. By holding the positive set constant and varying only the negative background, performance differences can be more directly attributed to background complexity rather than to shifts in the positive benchmark.

Beyond these primary benchmarks, we further assembled five additional validation datasets to examine specificity, sensitivity to rapidly emerging ARG families, and utility for large-scale metagenomic discovery (Fig. 2e).

### 2.3 GeoARG enhances detection of ARGs across sequence identities and input conditions

We evaluated GeoARG in two benchmark settings, GeoARG-DB-Test with UniProt [21] and GeoARG-DB-Test with NCBI open reading frames (ORFs), to compare performance under curated-protein and genome-derived backgrounds. In the UniProt benchmark (Fig. 3a,b), GeoARG achieved the strongest overall binary ARG identification performance among the tested methods, including an accuracy of 0.9992 and an MCC of 0.9984 on amino acid inputs (Supplementary Information Tables S2.1–S2.2).

**Fig. 3.**
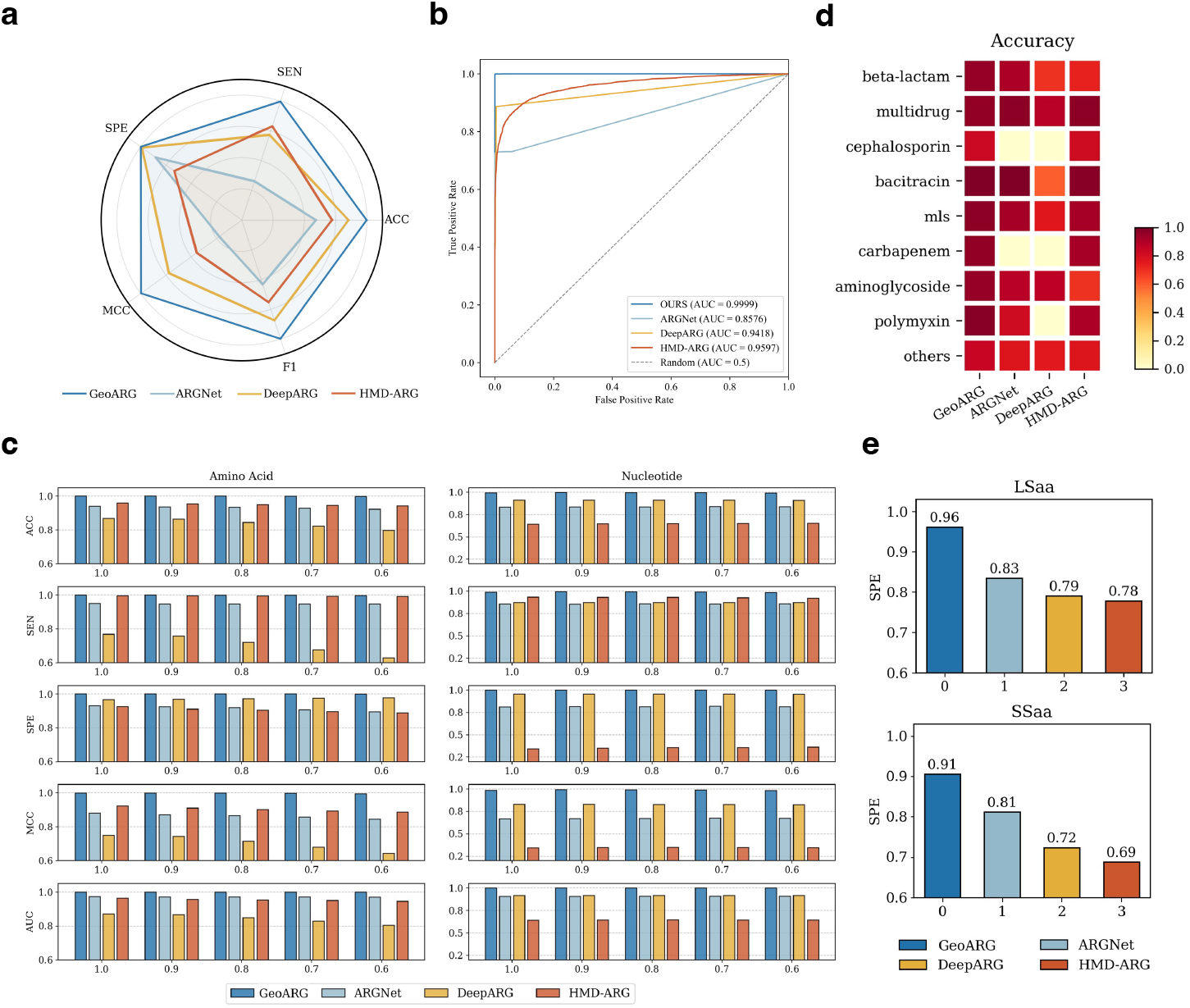
Performance benchmarking of GeoARG against state-of-the-art methods. **a** Radar chart on five classification metrics. **b** ROC curves on the UniProt benchmark. **c** Performance at decreasing sequence length thresholds (1.0–0.6). **d** Per-class accuracy heatmap across resistance categories. **e** Specificity against a virus protein dataset.

We then assessed robustness under the more complex ORF background (Fig. 3c), where negatives are less constrained and more similar to realistic metagenomic contexts. Across all four input settings (LSnt, SSnt, LSaa, and SSaa), GeoARG maintained high discrimination capacity, with MCC values above 0.88 and AUROC above 0.97. Performance remained stable as effective sequence length decreased, indicating that the model retains predictive signal even under short-fragment conditions.

Beyond binary prediction, we evaluated fine-grained annotation in a 36-class resistance-subtype task. GeoARG reached an overall accuracy of 0.8792, exceeding ARGNet (0.7380) and DeepARG (0.6260) (Fig. 3d). This gap suggests that gains are not limited to ARG/non-ARG separation, but also extend to subtype-level discrimination.

To analyze which components drive these improvements, we performed ablation experiments (Table 1). Removing structural features reduced MCC by 0.0509–0.1008 and AUROC by 0.0078– 0.1314, with larger decreases in short-sequence settings, where geometric information is expected to be more informative. Removing cross-attention also decreased performance (MCC drop: 0.0279– 0.1339; AUROC drop: 0.0014–0.0722), particularly in LSaa. Across all settings, the full model consistently outperformed ablated variants, supporting the joint contribution of structural encoding and multimodal fusion.

**Table 1.** Ablation study results across nucleotide and amino acid settings.

| Test setting | Model | Accuracy | Sensitivity | Specificity | MCC | AUROC |
| --- | --- | --- | --- | --- | --- | --- |
| LSnt | esm2_t12 fine-tuned | 0.9761 | 0.9654 | 0.9825 | 0.9187 | 0.9843 |
|  | w/o structure | 0.9879 | 0.9782 | 0.9937 | 0.9353 | 0.9911 |
|  | w/o cross attention | 0.9824 | 0.9853 | 0.9819 | 0.9583 | 0.9975 |
|  | **GeoARG** | **0.9931** | **0.9881** | **0.9983** | **0.9862** | **0.9989** |
| SSnt | esm2_t12 fine-tuned | 0.8194 | 0.8352 | 0.8098 | 0.7503 | 0.8417 |
|  | w/o structure | 0.8536 | 0.8677 | 0.8446 | 0.7963 | 0.8763 |
|  | w/o cross attention | 0.8974 | 0.8714 | 0.9099 | 0.8355 | 0.9104 |
|  | **GeoARG** | **0.9472** | **0.9131** | **0.9827** | **0.8969** | **0.9826** |
| LSaa | esm2_t12 fine-tuned | 0.9403 | 0.9471 | 0.9295 | 0.8724 | 0.9518 |
|  | w/o structure | 0.9590 | 0.9602 | 0.9484 | 0.9001 | 0.9672 |
|  | w/o cross attention | 0.8961 | 0.8789 | 0.9151 | 0.8634 | 0.9706 |
|  | **GeoARG** | **0.9987** | **0.9979** | **0.9994** | **0.9973** | **0.9998** |
| SSaa | esm2_t12 fine-tuned | 0.8071 | 0.7893 | 0.8196 | 0.7428 | 0.8189 |
|  | w/o structure | 0.8396 | 0.8221 | 0.8494 | 0.7887 | 0.8502 |
|  | w/o cross attention | 0.9141 | 0.8989 | 0.9257 | 0.8608 | 0.9373 |
|  | **GeoARG** | **0.9443** | **0.9220** | **0.9770** | **0.8895** | **0.9816** |

### 2.4 GeoARG maintains high specificity against evolutionarily distant non-bacterial sequence

Sequence-based ARG predictors risk elevated false-positive rates when applied to proteins that share superficial sequence features with ARGs but lack resistance function. To assess specificity under an extreme negative background, we evaluated GeoARG on a curated dataset of 8,793 viral proteins spanning 12 animal virus families from the NCBI RefSeq database [20]. Viral proteins are taxonomically remote from bacteria and carry no known antibiotic resistance functions, making this dataset a stringent test of false-positive control. GeoARG achieved high specificity on both long and short amino acid inputs (LSaa: 0.9611; SSaa: 0.9062), substantially outperforming ARGNet (LSaa: 0.8343; SSaa: 0.8124), HMD-ARG (LSaa: 0.7781; SSaa: 0.6883), and DeepARG (LSaa: 0.7900; SSaa: 0.7236) (Fig. 3e). The performance gap was consistent across both input lengths, suggesting that the structural constraints learned through knowledge distillation effectively suppress spurious predictions on proteins lacking resistance-associated catalytic architecture.

### 2.5 GeoARG exhibits biologically consistent prediction patterns on a low-ARG bacterial proteome

We evaluated genome-wide prediction behavior on the complete *E. coli* K12 reference proteome (Table 2). *E. coli* K12 is a laboratory-adapted strain with few known acquired ARGs and serves as a well-characterized reference background for assessing false-positive tendencies of ARG prediction models.

**Table 2.**
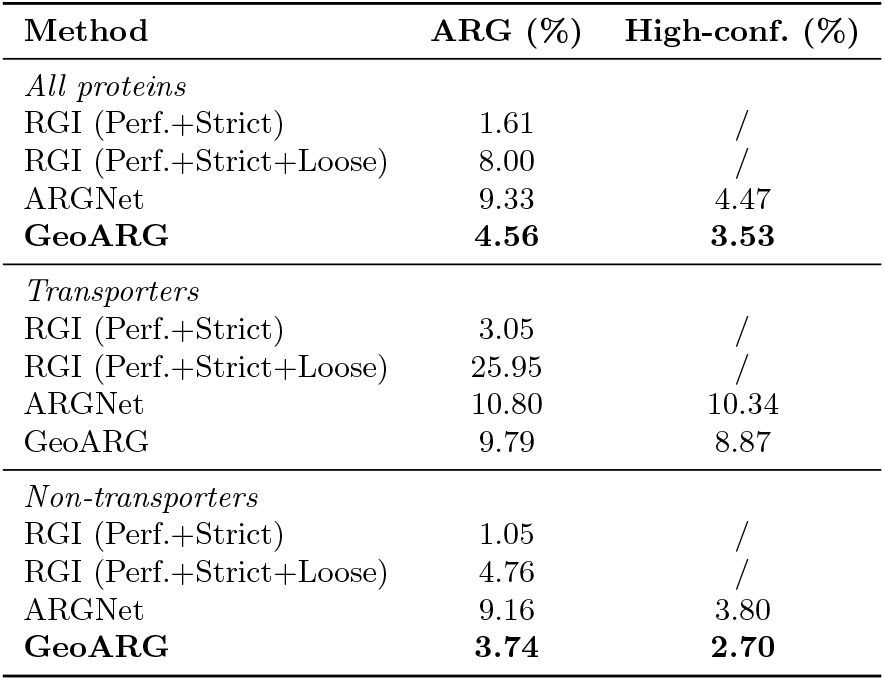
ARG prediction rates in the *E. coli* K12 proteome.

| Method | ARG (%) | High-conf. (%) |
| --- | --- | --- |
| *All proteins* |  |  |
| RGI (Perf.+Strict) | 1.61 | / |
| RGI (Perf.+Strict+Loose) | 8.00 | / |
| ARGNet | 9.33 | 4.47 |
| **GeoARG** | **4.56** | **3.53** |
| *Transporters* |  |  |
| RGI (Perf.+Strict) | 3.05 | / |
| RGI (Perf.+Strict+Loose) | 25.95 | / |
| ARGNet | 10.80 | 10.34 |
| GeoARG | 9.79 | 8.87 |
| *Non-transporters* |  |  |
| RGI (Perf.+Strict) | 1.05 | / |
| RGI (Perf.+Strict+Loose) | 4.76 | / |
| ARGNet | 9.16 | 3.80 |
| **GeoARG** | **3.74** | **2.70** |

**Table 3.** Recall on canonical and expanded *mcr* sequences.

| Setting | Model | Recall | Exp. recall |
| --- | --- | --- | --- |
| LSnt | ARGNet | **1.0000** | 0.9967 |
|  | **GeoARG** | **1.0000** | **0.9761** |
| SSnt | ARGNet | 0.8448 | 0.8506 |
|  | **GeoARG** | **0.9445** | **0.9569** |
| LSaa | ARGNet | 0.9991 | 0.6665 |
|  | **GeoARG** | **1.0000** | **0.9718** |
| SSaa | ARGNet | 0.7859 | 0.6281 |
|  | **GeoARG** | **0.9429** | **0.9532** |

At the whole-proteome level, GeoARG predicted 4.56% of proteins as ARGs (3.53% high-confidence), substantially lower than ARGNet (9.33%) and falling between RGI strict (1.61%) and RGI loose (8.00%) modes, suggesting a balanced sensitivity-specificity tradeoff without excessive false-positive predictions. Stratified analysis revealed a biologically consistent pattern. Among non-transporter proteins, which are less likely to harbor resistance-relevant functions, GeoARG predicted substantially fewer ARGs (3.74%) than ARGNet (9.16%), approaching RGI strict mode and indicating improved control of spurious predictions on non-resistance proteins. Among transporter proteins, where resistance-associated functions such as efflux-mediated drug export are more likely to reside, GeoARG maintained a prediction rate (9.79%) similar to ARGNet (10.80%) but markedly lower than RGI loose mode (25.95%). This stratified behavior—conservative on non-transporters, appropriately sensitive on transporters—suggests that GeoARG captures biologically relevant signals rather than uniformly suppressing predictions across all protein classes. We note that, in the absence of experimentally validated ARG labels for this proteome, these results reflect prediction tendencies rather than absolute accuracy.

### 2.6 GeoARG detects rapidly emerging resistance variants missed by reference databases

Public ARG databases such as CARD [13] are curated primarily from cultured pathogens, leaving newly emerged and phylogenetically divergent resistance variants systematically underrepresented. Mobilized colistin resistance (*mcr*) genes exemplify this challenge: they have diversified rapidly across gram-negative pathogens, generating a broad sequence landscape that extends well beyond canonical database entries. We therefore constructed validation sets spanning both canonical *mcr* genes and phylogenetically expanded *mcr*-like sequences, and assessed recall across four input settings (LSnt, SSnt, LSaa, SSaa).

GeoARG achieved perfect recall for canonical *mcr* sequences in both long-input settings (LSnt: 1.0000; LSaa: 1.0000), confirming sensitivity to well-characterized variants. More critically, GeoARG maintained high recall on phylogenetically expanded sequences where sequence divergence is greatest: on amino acid inputs, GeoARG substantially outperformed ARGNet (LSaa: 0.9718 vs. 0.6665; SSaa: 0.9532 vs. 0.6281), and advantages were consistent on short nucleotide inputs (SSnt: 0.9569 vs. 0.8506). ARGNet showed slightly higher expanded recall on LSnt (0.9967 vs. 0.9761), the setting where sequence context is most abundant and sequence-based methods are least disadvantaged. The performance gap widened markedly as sequence length decreased, consistent with geometric features providing discriminative signal precisely where sequential context is insufficient.

These results suggest that structural geometry captures functional conservation that persists across the evolutionary diversification of *mcr* variants — a property that sequence-based methods cannot reliably exploit, particularly for the truncated or divergent inputs typical of metagenomic data.

### 2.7 Discovery of remote ARG candidates from metagenomic datasets

Independent metagenomic validation was performed using ResFinderFG v2.0 [22] and The Global Microbial Gene Catalog (GMGC) gut catalog [23].

ResFinderFG v2.0 contains experimentally validated ARGs and is independent of GeoARG-DB and other public ARG resources, which is suitable for GeoARG’s generalization testing. Since all sequences in ResFinderFG v2.0 are experimentally validated ARGs, this dataset evaluates sensitivity exclusively; recall is therefore the primary metric reported. We grouped sequences by identity to GeoARG-DB into three bins (0–40%, 40–60%, and 60–100%); results are shown in Fig. 4a. In the 0–40% identity bin, GeoARG achieved 0.48 recall, compared with 0.11 for HMD-ARG and 0.02 for ARGNet. The advantage remained in the 40–60% bin (0.88 vs. 0.24 and 0.18), while all methods performed well in the 60–100% bin. These results indicate improved remote homolog detection by GeoARG (Supplementary Information Table S1.5).

**Fig. 4.**
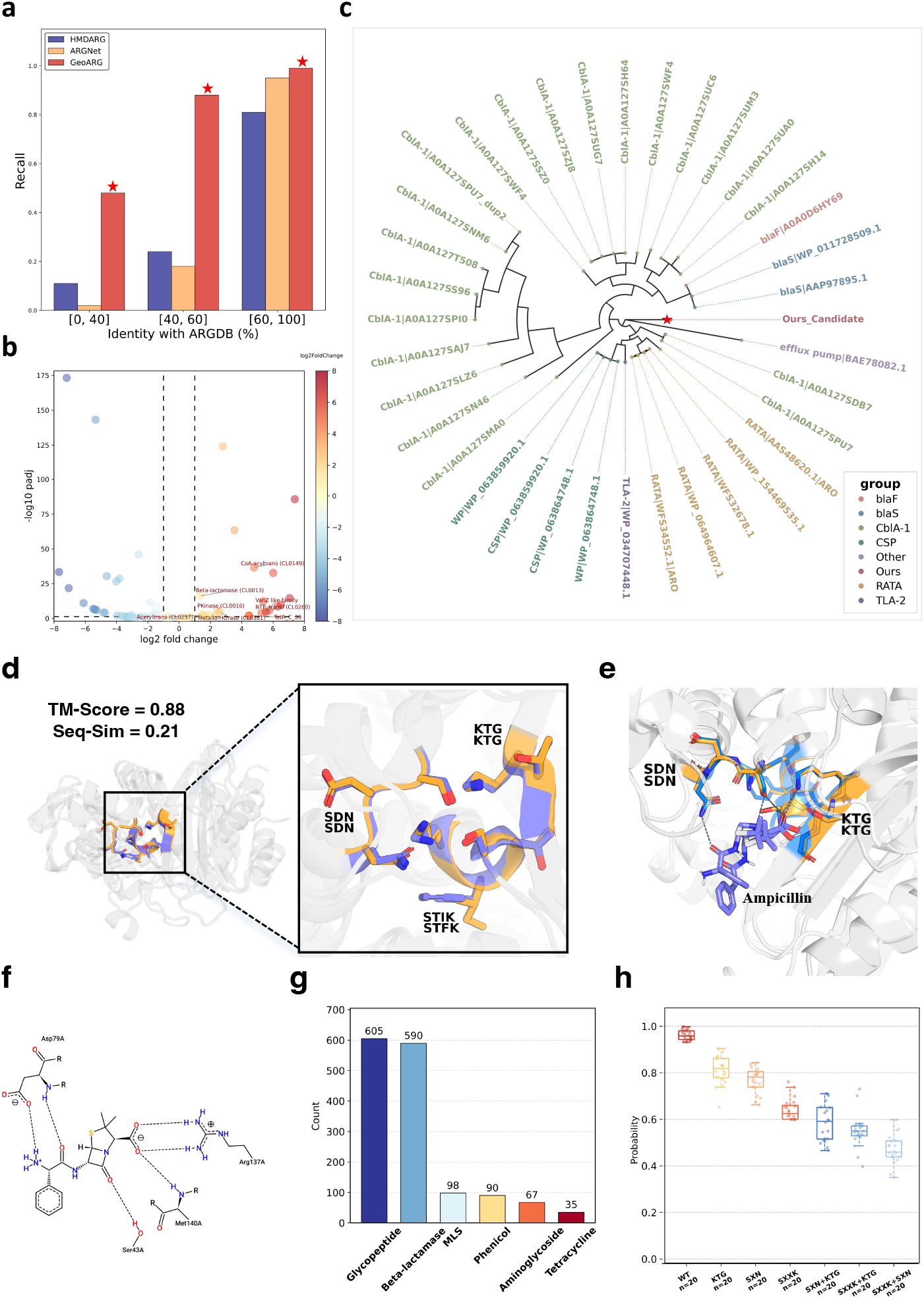
Metagenomic validation and structural analysis of novel ARG candidates. **a** Recall rates across identity bins. **b**,**c** Pfam enrichment volcano plot and phylogenetic candidate placement. **d** Structural alignment. **e**,**f** Predicted ligand binding. **g** GeoARG’s high-confidence candidates distributions. **h** Impact of in silico motif perturbations.

We next evaluated GeoARG on the GMGC gut dataset [23], a large-scale human gut metagenomic resource comprising over 52 million predicted ORFs. To focus on evolutionarily remote sequences, we filtered the dataset to retain only those sharing less than 25% sequence identity to any entry in GeoARG-DB, yielding 111,202 candidate sequences. Among these, 49,707 carried “UNKNOWN” annotation in the GMGC metadata, and analysis was restricted to this subset to focus on novel ARG discovery (Supplementary Information Section S1.8).

GeoARG was applied to these 49,707 sequences. Predictions annotated as “multidrug” or “unclassified” were excluded, as these categories lack a defined mechanistic target and are not amenable to downstream validation. Among the remaining predictions, GeoARG identified 1,485 high-confidence ARG candidates (*P >* 0.8), spanning six resistance classes: glycopeptide (605), beta-lactamase (590), MLS (98), phenicol (90), aminoglycoside (67), and tetracycline (35).

To validate the functional relevance of these candidates, we performed Pfam domain enrichment analysis on the 1,485-candidate set (Fig. 4b), comparing against 3,000 low-confidence background sequences (*P <* 0.2) drawn from the same unannotated GMGC sequence space (Supplementary Information Section S6). Of 148 Pfam features tested, 36 were significantly enriched and 66 depleted (FDR *<* 0.05; Benjamini–Hochberg correction), far exceeding random expectation (permutation mean = 0.08). Among the enriched features, multiple canonical resistance-associated domain families were identified, including the chloramphenicol acetyltransferase (CAT) family, VanS-type histidine kinase domains (HisKA, HATPase c), beta-lactamase folds (Beta-lactamase2, Lactamase B), and aminoglycoside-modifying enzyme domains (APH, NTP transf 2), collectively consistent with the predicted resistance-class composition of the candidate set.

### 2.8 High-confidence candidates are evolutionarily divergent yet functionally conserved

We first assessed sequence-level divergence by searching the 1,485 high-confidence candidates against the NCBI non-redundant database using DIAMOND (Fig. 5a). Across all resistance classes, a substantial fraction of sequences yielded no detectable genus-level hits, indicating little similarity to any currently annotated protein. This pattern was most pronounced for phenicol, aminoglycoside, and MLS candidates, confirming that these sequences are largely invisible to standard similarity-based detection (Supplementary Information Sections S6.1–S6.2).

**Fig. 5.**
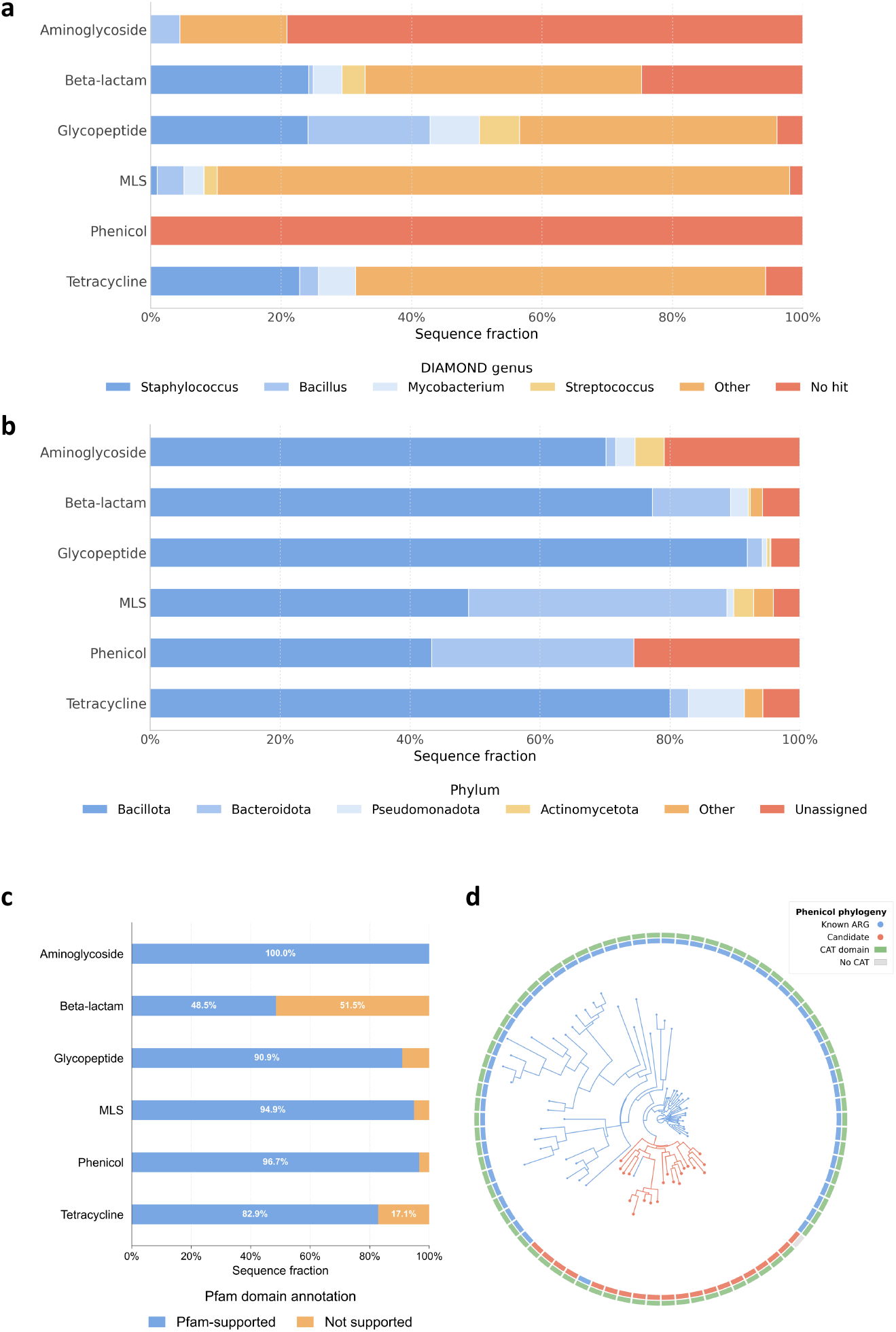
Evolutionary characterization of predicted ARG candidates. **a** DIAMOND genus-level best-hit distribution. **b** Phylum-level taxonomic distribution. **c** Pfam domain annotation rates across resistance classes. **d** Circular phylogeny of phenicol sequences (58 known, 20 candidates; clustered at 50% identity). Inner ring: sequence type. Outer ring: CAT domain annotation.

Despite extensive sequence divergence, the taxonomic distribution of candidates was markedly non-random (Fig. 5b). The majority were assigned to Bacillota, with additional contributions from Bacteroidota across multiple resistance classes—both dominant phyla in the human gut microbiome and well-established reservoirs of ARGs. This phylum-level concentration within ecologically and microbiologically relevant lineages is consistent with a biologically meaningful signal rather than random model noise.

Functional conservation was nonetheless detectable at the domain level (Fig. 5c). For most resistance classes, candidates carried canonical resistance-associated Pfam domains at high rates: aminoglycoside candidates were fully supported (100.0%), with strong annotation rates for MLS (94.9%), glycopeptide (90.9%), and tetracycline (82.9%). The most striking contrast emerged for phenicol: despite being almost entirely undetectable by sequence similarity search, 96.7% of phenicol candidates retained chloramphenicol acetyltransferase (CAT) domain annotation. Phylogenetic reconstruction confirmed this pattern (Fig. 5d): predicted candidates formed multiple distinct clades separated from known ARGs, yet CAT domain annotation was near-universally retained across these phylogenetically remote sequences—directly demonstrating that functional architecture is preserved even as sequences diverge beyond homology detection thresholds.

Beta-lactamase candidates were a notable exception, with only 48.47% carrying detectable Pfam annotation. However, 68.14% retained at least two of the three canonical serine beta-lactamase catalytic motifs (SXXK, SXN, and KTG; Table 4), suggesting that these candidates preserve the residues most critical for catalysis despite sufficient global sequence divergence to fall below Pfam profile HMM detection thresholds. This discrepancy motivated further structure and motif-level investigation.

**Table 4.** Motif conservation and Pfam annotation among high-confidence beta-lactamase candidates.

| Model | $N$ ($P > 0.8$) | SXXK (%) | SXN (%) | KTG (%) | $\geq 2$ motifs (%) | Pfam beta-lactamase (%) |
| --- | --- | --- | --- | --- | --- | --- |
| GeoARG | 590 | 79.15 | 75.93 | 52.00 | 68.14 | 48.47 |

### 2.9 Structural analysis reveals conserved catalytic architecture in remotely homologous ARGs

Beta-lactamase catalytic activity depends critically on the three-dimensional arrangement of active-site residues: the canonical SXXK, SXN, and KTG motifs must be correctly positioned in space to coordinate the catalytic serine and mediate substrate binding. Sequence-level divergence therefore does not preclude functional conservation if the underlying three-dimensional architecture is maintained. To assess whether high-confidence candidates preserve this geometric constraint despite low sequence identity, we performed structural characterization of representative candidates across two resistance classes.

The candidate `GMGC10.207_616_678.UNKNOWN` was selected on the basis of high prediction confidence (*P >* 0.99) and complete retention of all three canonical catalytic motifs. It shares only 21.11% sequence identity with its closest annotated beta-lactamase. Standard annotation pipelines failed to recognize its resistance function: NCBI-BLAST [9] returned only a generic “serine hydrolase” annotation, and the candidate was not detected by ARGNet or RGI (loose mode). This case exemplifies precisely the failure mode that motivated GeoARG: a functionally plausible resistance gene systematically missed by both sequence similarity search and existing deep learning tools.

Despite low global sequence similarity, the candidate exhibited high structural prediction confidence (pLDDT = 96.27) and retained all three canonical catalytic motifs of class A beta-lactamases (Fig. 4d; Supplementary Information Section S4). Phylogenetic analysis placed the candidate within the class A beta-lactamase clade rather than with unrelated serine hydrolases (Fig. 4c), suggesting that GeoARG captures lineage-related functional features rather than superficial sequence similarity. Foldseek [24] structural alignment with a reference class A beta-lactamase (`WP_064935627.1`) yielded a TM-score of 0.88 and a motif-level C*α* RMSD of 0.32 Å. AlphaFold3 co-folding of ampicillin with both the candidate and reference protein produced highly similar binding poses (motif-level C*α* RMSD = 0.29 Å; Fig. 4e–f), with ampicillin localizing to the canonical catalytic pocket and forming hydrogen bonds with residues within the conserved motifs.

Structural conservation of catalytic architecture was not limited to the beta-lactamase class. A representative glycopeptide candidate, `GMGC10_052_200_458.UNKNOWN`, shares only 14.1% sequence identity with VanS-D (`WP_063856738.1`), yet Foldseek alignment yielded a TM-score of 0.85 and pairwise C*α* distances of 1.0–2.7 Å across the conserved H-box, N-box, and G-box motifs, confirming strong spatial conservation of the catalytic architecture despite extreme sequence divergence.

### 2.10 Motif-level perturbation analysis reveals catalytic residue sensitivity

To interpret which input features drive GeoARG predictions, we performed counterfactual perturbation analysis on 20 high-confidence beta-lactamase candidates (*P >* 0.80, pLDDT *>* 90, all three canonical motifs retained). Canonical catalytic-motif residues were substituted with alanine in silico under six perturbation conditions: individual disruption of each motif (KTG, SXN, SXXK) and pair-wise disruption of any two motifs simultaneously (SXN+KTG, SXXK+KTG, SXXK+SXN). Each condition was applied independently to all 20 sequences, yielding *n* = 20 per group. The resulting sequences were passed through the GeoARG inference pipeline and predicted probabilities were compared against unperturbed wild-type sequences (WT, median = 0.96; Fig. 4h).

Among single-motif disruptions, the three motifs contributed unequally to model confidence. Disruption of SXXK produced the largest probability reduction (median = 0.65), followed by SXN (median = 0.78) and KTG (median = 0.81), suggesting that the SXXK motif, which contains the nucleophilic serine required for beta-lactamase activity, is the primary determinant of GeoARG’s resistance predictions. Pairwise disruption consistently amplified these reductions: SXN+KTG (median = 0.59), SXXK+KTG (median = 0.55), and SXXK+SXN (median = 0.48) all fell substantially below any single-motif condition, indicating that the three motifs contribute independently and additively to model confidence. The ordering of Combo conditions further reflects the dominant role of SXXK: pairwise combinations involving SXXK (SXXK+KTG, SXXK+SXN) produced greater reductions than the combination excluding it (SXN+KTG).

These results indicate that GeoARG predictions are sensitive to the integrity of catalytic residues in a manner consistent with known beta-lactamase biochemistry, where the SXXK motif anchors the active-site serine and coordinates substrate acylation. The graded, motif-specific response across all 20 sequences suggests that this sensitivity is a systematic property of the model rather than an artifact of individual sequence context, supporting the interpretation that GeoARG captures functional signals at the resolution of individual catalytic residues.

### 2.11 Molecular dynamics analysis supports stability of the predicted ligand binding

While structural alignment and docking support the plausibility of ligand binding, these analyses remain inherently static. To further assess whether the predicted interaction is dynamically stable under physiological conditions, we performed molecular dynamics (MD) simulations. We simulated the AlphaFold3-predicted complex between the candidate protein and ampicillin. The ligand remained positioned within the catalytic pocket throughout the simulation, maintaining interactions with residues surrounding the conserved active-site motifs (Fig. 6a).

**Fig. 6.**
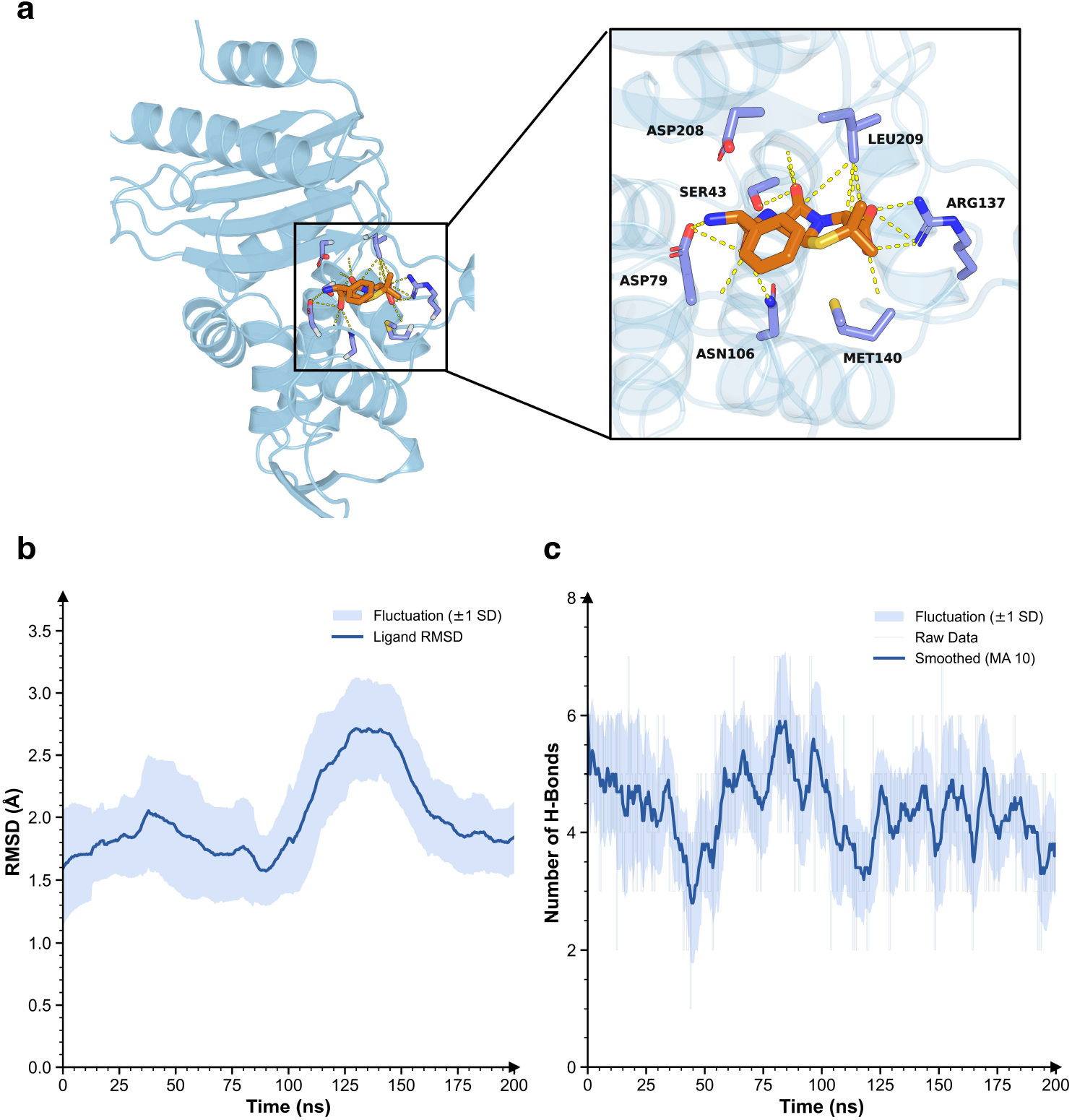
Molecular dynamics of the predicted protein-ligand complex. **a** Binding pose of ampicillin. **b** Ligand RMSD trajectory over 200 ns. **c** Hydrogen bond fluctuations during simulation.

The stability of the ligand-binding configuration was assessed using ligand root-mean-square deviation (RMSD) relative to the initial structure. As shown in Fig. 6b, the ligand RMSD fluctuated within approximately 1–3 Å during the simulation and did not exhibit progressive drift over time, indicating that the predicted binding pose remained structurally stable within the catalytic pocket. Consistent with this observation, hydrogen bond analysis revealed persistent interactions between ampicillin and surrounding residues (Fig. 6c), typically ranging between three and six interactions involving residues located near the conserved catalytic motifs.

Together, these results indicate that the predicted protein–ligand complex remains dynamically stable during MD simulation and preserves interactions consistent with the catalytic environment of class A beta-lactamases. Although biochemical validation is required to confirm enzymatic activity, these structural and dynamical observations further support the functional plausibility of the GeoARG-predicted ARG candidate (Supplementary Information Section S7).

### 2.12 Computational efficiency of knowledge distillation

Inference efficiency was evaluated on the GeoARG-DB-Test dataset (16,250 protein sequences) using a single NVIDIA A100 GPU with a batch size of 16. Runtime and parameter size were compared between the teacher model (sequence + structure encoders) and the distilled student model. The student model achieved substantially faster inference while maintaining prediction performance (Table 5).

**Table 5.** Inference efficiency comparison between the teacher and student models.

| Model | Params | Time (s) | Speedup |
| --- | --- | --- | --- |
| Teacher (ESM2-650M + E(3) GNN) | 665.25M | 1709 | 1× |
| Student (ESM2-35M) | 33.77M | 111 | 15.4× |

## 3 Discussion

PLMs have significantly improved protein sequence representation by capturing evolutionary patterns from large-scale datasets [25–27]. PLMs’ representations enable models to learn long-range dependencies within sequences and have shown competitive performance in protein function prediction tasks [25, 28, 29]. However, sequence-based representations including PLMs and alignment-based methods, are insufficient for identifying ARGs that have undergone large sequence divergence, since ARG function is often determined by the three-dimensional geometry of catalytic residues [17, 18]. Sequence-based methods therefore fail to detect ARGs that have diverged in sequence but retained catalytic function, limiting the discovery of novel resistance genes.

GeoARG addresses this challenge by explicitly modeling the three-dimensional structure of residues through an E(3)-equivariant graph neural network [30], which encodes pairwise geometric relationships between backbone atoms in a manner invariant to global rotation and translation. The ablation results support this: removing structural features caused the largest performance drops in short-sequence settings (SSnt/SSaa; MCC drop up to 0.1008, AUROC up to 0.1314), where limited sequential context makes geometric signals most decisive. These design choices allow GeoARG to capture conserved catalytic architecture even when global sequence identity falls well below conventional homology thresholds.

A practical challenge in applying structure-aware models to large-scale metagenomic data is computational cost. The knowledge distillation framework addresses this directly: by transferring multimodal representations into a lightweight student PLM (33.77M parameters), GeoARG achieves a 15.4-fold speedup while retaining predictive performance, making the analysis of large datasets such as the GMGC gut catalog [23] tractable in practice.

Large-scale metagenomic sequencing has revealed that the majority of environmental protein-coding sequences share little similarity with curated reference databases [31]. Our analysis of 49,707 unannotated GMGC gut sequences identified 1,485 high-confidence ARG candidates, with Pfam enrichment analysis confirming significant overrepresentation of canonical resistance-associated families (e.g., CAT family: OR = 240.07, FDR = 4.6 × 10^−32^). The beta-lactamase case study further illustrates that structure-informed prediction can recover functionally specific annotations from sequences that alignment-based pipelines systematically misclassify or leave unannotated.

Several limitations merit acknowledgment. Protein structures were predicted by ESMFold rather than resolved experimentally. Although ESMFold generally yields high-confidence predictions for known protein families (median pLDDT = 94.9), and the representative candidate analyzed here achieved a pLDDT of 96.27, the influence of structural prediction uncertainty on GNN representations in lower-confidence cases remains uncharacterized. Most importantly, molecular dynamics simulations support physical plausibility but cannot replace experiments.

Overall, this study demonstrates that integrating geometric structural constraints with PLMs substantially improves the detection of remotely homologous ARGs and enables systematic discovery of novel resistance candidates in large-scale metagenomic datasets. These findings suggest that structure-informed representation learning offers a broadly applicable strategy for expanding the known resistome and may inform future AMR surveillance in complex microbial communities.

## 4 Methods

### 4.1 Curation and annotation of GeoARG-DB

GeoARG-DB was constructed by integrating experimentally validated ARG protein sequences from seven publicly available resources, including ARGNet-DB [15], HMD-ARG-DB [16], DeepARG-DB [14], ResFinder [32], AMRFinder [33], MEGARes [34–36], and CARD [11–13]. Merging all sources yielded 58,771 amino acid sequences. Entries annotated as resistance determinants solely through single-nucleotide polymorphisms were excluded as they do not represent transferable ARG protein families. Redundancy was then removed using CD-HIT (v4.8.1) [37] at 100% sequence identity and full-length coverage, resulting in a curated set of 40,624 non-redundant ARG proteins (Fig. 2a).

All sequences were then mapped to a standardized resistance annotation framework and categorized into 36 resistance classes. Proteins associated with multiple resistance mechanisms were assigned a “multidrug” label. This unified annotation system was applied consistently across model training, subtype evaluation, and all downstream analyses (Supplementary Information Section S3).

### 4.2 Benchmark setup and negative background generation

GeoARG-DB proteins were split at the sequence level into training (GeoARG-DB-Train, 70%), validation (GeoARG-DB-Val, 10%), and test (GeoARG-DB-Test, 20%) sets. For both model training and validation, negative sets were subsampled to maintain class balance with the corresponding positives in each split. To construct a proteome-style negative background, bacterial proteins were sampled from UniProt [21] across 1,000 species. To reduce sequence-length bias, negatives were selected to match the length distribution of GeoARG-DB positives. Proteins annotated with resistance-related functions were excluded, and sequences from *Escherichia coli* (*E. coli*) sequences were removed to avoid overlap with downstream proteome-level evaluations.

For the genome-style negative background, 1,000 bacterial reference genomes were obtained from NCBI (June 2025) [20], and open reading frames (ORFs) were predicted using Prokka [38]. Candidate negatives from both sources were aligned to GeoARG-DB using BLASTx [9], and sequences meeting homology thresholds (E-value ≤ 1 × 10^−5^, identity ≥ 20%, and coverage ≥ 70%) were removed (Fig. 2c). Two held-out evaluation benchmarks were defined by pairing the same positive set (GeoARG-DB-Test) with distinct negative backgrounds: (i) UniProt-derived bacterial proteins and (ii) genome-derived ORFs. The UniProt benchmark consisted of full-length amino acid sequences, whereas the ORF benchmark included both full-length and truncated inputs for LSaa and LSnt evaluation. Formal definitions of LSnt, SSnt, LSaa, and SSaa are described below.

### 4.3 External validation datasets

#### 4.3.1 Virus dataset

Viral protein sequences were incorporated as an additional negative test dataset. A total of 8,793 protein sequences from 12 animal virus families were downloaded from the NCBI Protein RefSeq database [20] (accession date: 12 November 2025). The dataset comprised sequences from Arenaviridae (247), Coronaviridae (1,098), Filoviridae (127), Flaviviridae (1,076), Hantaviridae (151), Orthomyxoviridae (241), Paramyxoviridae (699), Phenuiviridae (609), Picornaviridae (1,229), Pneumoviridae (106), Reoviridae (1,094), and Rhabdoviridae (2,116).

#### 4.3.2 *E. coli* K-12 dataset

To evaluate model behavior on a well-annotated bacterial background, 66 complete *E. coli* K-12 genomes were obtained from NCBI [20] (accession date: 18 December 2025). Following redundancy removal, a non-redundant set of 4,300 protein sequences was retained as the test dataset. Based on NCBI functional annotations, 435 proteins were labeled as transporters.

#### 4.3.3 *mcr*-like dataset

To construct an expanded *mcr*-like evaluation dataset, two distantly related RefSeq proteins, *mcr-1* (`WP_163397051.1`) and *mcr*-4 (`WP_099156046.1`), were used as query sequences in tBLASTn [9] searches against the NCBI nucleotide database. For each query, the top 5,000 hits ranked by ascending E-value were retained. Corresponding GenBank records was retrieved, and all coding sequences (CDS) features overlapping the aligned regions were extracted to obtain protein sequences for down-stream analysis. An additional 17,071 *mcr*-labelled genes were collected from the NCBI MicroBIGG-E web service (https://www.ncbi.nlm.nih.gov/pathogens/microbigge/). We further incorporated 324 EptA/B/C (*mcr*-like) sequences reported in a comparative study [39], with no overlap with other sources. After merging and removing duplicates, a total of 29,352 non-redundant protein sequences were retained.

All sequence alignments were then performed using MAFFT (FFT-NS-i) [40], and a maximum-likelihood phylogenetic tree was constructed using FastTree under the LG+CAT model [41]. Based on source metadata, 17,071 sequences were initially labeled as “*mcr* “. To expand this definition, the most recent common ancestor (MRCA) of each *mcr* subtype (*mcr*-1 to *mcr*-10) was identified within the inferred tree, and all descendant sequences were inccluded in the expanded *mcr* set, following Pei et al. [15]. This procedure resulted in a total of 19,163 expanded *mcr* sequences.

#### 4.3.4 ResFinderFG and GMGC datasets

Experimentally validated ARGs from ResFinderFG v2.0 [22] were used as an independent benchmark. Sequence identities between ResFinderFG proteins and GeoARG-DB were computed using DIAMOND blastp (v2.1) [42] with an e-value threshold of 1 × 10^−5^, and proteins were grouped into 0–40%, 40–60%, and 60–100% identity bins.

For large-scale discovery, GMGC gut proteins [23] were aligned to GeoARG-DB using DIAMOND [10]. Proteins sharing ≥ 25% sequence identity were removed, and entries annotated as “unknown protein” were retained, resulting in 49,707 sequences.

### 4.4 GeoARG architecture

GeoARG follows a teacher–student architecture with three functional components: sequence/structure feature encoding, multimodal fusion with distillation, and lightweight prediction. The teacher comprises a pretrained PLM (`esm2_t33_650M_UR50D`) [25] and an E(3)-equivariant graph neural network [30]. The student is a smaller PLM (`esm2_t12_35M_UR50D`) that receives sequence input only at inference.

During training, sequence and structure features are jointly modeled in the teacher and transferred to the student through distillation. During inference, only the distilled student is retained, enabling efficient large-scale ARG screening.

#### 4.4.1 Protein structure prediction and geometric graph construction

Protein three-dimensional structures were predicted from amino acid sequences using ESMFold [25]. Per-residue confidence was assessed using pLDDT scores; residues with pLDDT *<* 70 were excluded during geometric feature construction.

Predicted structures were represented as residue-level graphs 𝒢= (𝒱, ℰ), where each residue corresponds to a node *i* ∈ 𝒱 with backbone coordinate *x*_*i*_ ∈ ℝ^3^. An edge (*i, j*) ∈ ℰ was constructed if the C*α*–C*α* distance *d*_*ij*_ = ∥*x*_*i*_ − *x*_*j*_∥_2_ was below 8 Å. Node features concatenated a 20-dimensional one-hot encoding of amino acid identity, backbone dihedral angles (*ϕ, ψ, ω*) encoded as sine–cosine pairs (6 dimensions), and the per-residue pLDDT score as a scalar confidence weight, giving a total node feature dimension of 27.

Residue graphs were processed by an E(3)-equivariant graph neural network [30] with *L* = 6 message-passing layers and hidden dimension *d*_*h*_ = 512. Learnable functions were implemented as two-layer MLPs with SiLU activations and layer normalisation, with residual connections at each layer.

The graph encoder satisfies full E(3) equivariance — invariance to rotations, translations, and reflections — through two design choices. First, edge messages depend only on the squared pairwise distance and pLDDT confidence,

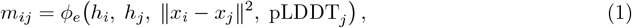

which are scalar quantities invariant under any orthogonal transformation of the coordinate frame. Second, *C*_*α*_ coordinates are updated via a weighted sum of unit radial directions,

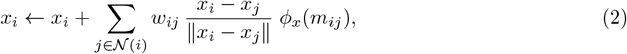

where *ϕ*_*x*_ is a scalar-valued MLP and *w*_*ij*_ are pLDDT-normalised aggregation weights,

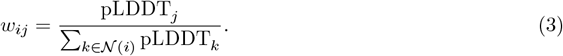

Because the direction vector (*x*_*i*_−*x*_*j*_)*/*∥*x*_*i*_−*x*_*j*_∥ transforms covariantly under any rotation or reflection applied to the input coordinates, and *ϕ*_*x*_(*m*_*ij*_) is a rotation-invariant scalar, the coordinate update is E(3)-equivariant by construction. Node features *h*_*i*_ are updated using only aggregated scalar messages, preserving their E(3)-invariance throughout all layers. A global mean-pooling readout produced the structure embedding *z*_struc_ ∈ ℝ^512^.

#### 4.4.2 Multimodal fusion and knowledge distillation

Sequence and structure representations were fused through a cross-attention module [43], where sequence features serve as queries and structure features as keys/values:

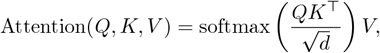

where *d* is the embedding dimension. The fused teacher representation was

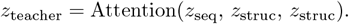

Knowledge distillation [44] was applied to transfer multimodal knowledge to the student with objective

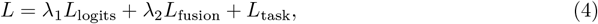

where *L*_task_ is task cross-entropy. The logits loss *L*_logits_ was defined as temperature-scaled KL divergence:

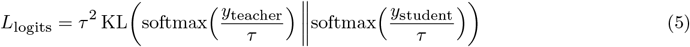

where *τ* is the temperature parameter. The fusion alignment loss *L*_fusion_ minimized the mean squared error between the fused teacher embedding *z*_teacher_ and the student embedding *z*_student_:

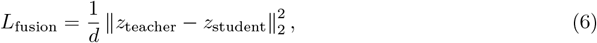

where *d* is the embedding dimension.

### 4.5 Training procedure and hyperparameters

All teacher components were frozen, and only the student model plus prediction head were optimized. The prediction head produced logits for binary ARG identification and subtype classification. Optimization used AdamW with weight decay [45], and early stopping was applied using validation performance (Supplementary Information Section S1.1–S1.2).

Training used GeoARG-DB-Train, hyperparameter tuning and early stopping used GeoARG-DB-Val, and final evaluation used the independent test set only. Models were implemented in PyTorch [46] and trained on a single NVIDIA A100 GPU. Core training settings included teacher-frozen/student-only optimization and distillation with (*λ*_1_, *λ*_2_, *τ*) as defined above; detailed numeric hyperparameters are provided in the Supplementary Information (Section S1.1–S1.2).

### 4.6 Baselines and unified comparison rules

To enable fair comparison across methods with heterogeneous outputs (GeoARG, ARGNet, DeepARG, HMD-ARG, and RGI), unified binary rules were applied. For DeepARG, a sequence was considered positive if any hit was returned; no-hit sequences were treated as negative. For RGI, predictions labeled Perfect, Strict, or Loose were each evaluated as positive calls in separate analyses.

For deep learning models producing probabilities (GeoARG, ARGNet, HMD-ARG), binary classification used *P >* 0.5. High-confidence analyses used a stricter threshold of *P >* 0.8, consistent with prior ARGNet settings [15]. DRAMMA was excluded from direct comparison because it requires nucleotide contig context, which is incompatible with the protein-sequence-only comparison setting used here.

#### 4.6.1 Nucleotide sequence processing

For nucleotide input settings (LSnt and SSnt), input sequences were translated to amino acid sequences using six-frame translation implemented with Biopython [47]. All six reading frames (three forward, three reverse-complement) were translated, and the frame yielding the longest open reading frame — defined as the longest stretch between stop codons — was retained. Translated sequences were then processed identically to amino acid inputs through the student PLM pipeline.

### 4.7 Structural and downstream analyses

For functional enrichment of predicted candidates, Pfam enrichment analysis was performed using HMMER (v3.3) [48, 49] against Pfam-A [50, 51] with an E-value threshold of 1 × 10^−5^. Detected domains were mapped to Pfam clans and merged within clans before enrichment testing.

For structural plausibility analyses of representative remote-homology candidates, structures were compared using Foldseek [24] and motif-level geometric consistency was assessed by C*α*-based structural comparisons around catalytic motifs. Counterfactual motif perturbations were performed by substituting canonical catalytic-motif residues with alanine across 20 high-confidence candidates.

For ligand-level structural assessment, AlphaFold3 co-folding [52] was used to predict protein– ligand complexes for selected candidates and references. Molecular dynamics simulations were then conducted in GROMACS [53] with CHARMM36m [54] and TIP3P water [55], and trajectories were analyzed in PyMOL [56].

### 4.8 Evaluation metrics and statistical analysis

Model performance was evaluated using accuracy, precision, recall, Matthews correlation coefficient (MCC), area under the receiver operating characteristic curve (AUROC), and where applicable area under the precision-recall curve (AUPRC). These metrics were computed from true positives (TP), true negatives (TN), false positives (FP), and false negatives (FN) as follows:

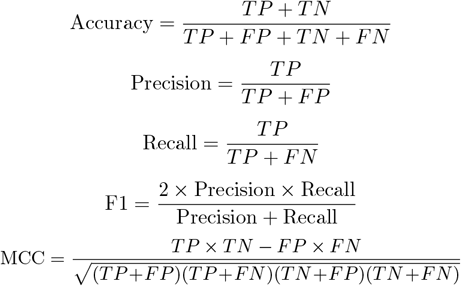

MCC provides a balanced measure of binary classification performance that is robust to class imbalance, making it particularly suitable for ARG identification tasks where positive and negative samples may differ substantially in number. AUROC summarizes discrimination ability across all decision thresholds, while AUPRC is additionally reported in settings where the positive class is rare.

For the virus specificity evaluation, identification specificity was defined as the proportion of viral sequences correctly classified as non-ARG. For the *E. coli* K12 proteome analysis, ARG prediction rate was computed separately for transporter and non-transporter subsets based on NCBI functional annotations.

For enrichment analyses, Fisher’s exact test was applied on 2 × 2 contingency tables, odds ratios were computed with a pseudocount of 0.5 to avoid undefined values, and multiple testing correction used the Benjamini–Hochberg procedure [57]. Features with FDR *<* 0.05 were considered statistically significant.

### 4.9 Web server availability

GeoARG is freely available through a public web server at https://ycclab.cuhk.edu.cn/GeoARG/, which supports online ARG prediction for user-submitted sequences.

## Data availability

Data is available at https://github.com/XingqiaoLin/GeoARG and https://zenodo.org/records/19296092. GeoARG-DB was constructed from seven public ARG resources: ARGNet-DB, HMD-ARG-DB, DeepARG-DB, ResFinder, AMRFinder, MEGARes, and CARD. The UniProt bacterial protein dataset is available at https://www.uniprot.org. Bacterial reference genomes and viral protein sequences were obtained from NCBI RefSeq at https://www.ncbi.nlm.nih.gov. The mcrlabelled genes were collected from the NCBI MicroBIGG-E web service at https://www.ncbi.nlm.nih.gov/pathogens/microbigge/. ResFinderFG v2.0 is available at https://github.com/RemiGSC/ResFinder_FG_Construction. The Global Microbial Gene Catalog (GMGC) gut dataset is available at https://gmgc.embl.de.

## Code availability

Code is available at https://github.com/XingqiaoLin/GeoARG, including trained weights, inference scripts, and training scripts. Trained model weights are also available at https://zenodo.org/records/19296092. The GeoARG web server is freely accessible at https://ycclab.cuhk.edu.cn/GeoARG/. Structural alignments used Foldseek (https://github.com/steineggerlab/foldseek), protein–ligand co-folding used AlphaFold3 (https://github.com/google-deepmind/alphafold3), and structure visualizations used PyMOL (https://pymol.org).

## Acknowledgements

This work was supported by the Fundamental Research Funds for the Central Universities (20720250172).This work was supported by the Scientific Research Foundation of State Key Laboratory of Vaccines for Infectious Diseases, Xiang An Biomedicine Laboratory(2025XAKJ0102017).

## Conflict of interest

The authors declare no conflict of interest.

## Supplementary information

Supplementary information is available for this paper.

## Notes

https://github.com/XingqiaoLin/GeoARG

https://zenodo.org/records/19296092

https://ycclab.cuhk.edu.cn/GeoARG/

### Competing Interest Statement

The authors have declared no competing interest.

