## Supplementary Information for "Geometry-enhanced protein language modeling enables discovery of novel antibiotic resistance genes"

### GeoARG SI

#### S1. Hyperparameters and Training Details

##### S1.1 Model Architecture

Table 1: E(3)-equivariant GNN hyperparameters

| Hyperparameter | Value |
|---|---|
| Message-passing layers | 6 |
| Hidden / output dimension | 512 |
| $C_\alpha$–$C_\alpha$ edge cutoff | 8.0 Å |
| Node feature dimension | 27 |
| Edge message input | $[h_i \| h_j \| \|x_i - x_j\|^2 \| \mathrm{pLDDT}_j]$ |
| Activation | SiLU |
| Normalisation | LayerNorm + residual |
| Coordinate update | Radial direction × scalar weight (E(3)-equivariant) |
| Symmetry group | E(3) (rotation, translation, reflection) |

Table 2: Teacher model parameter counts

| Module | Total params | Trainable params |
|---|---|---|
| ESM2-650M (backbone) | 651.04M | 78.71M |
| ESM2 projection | 0.66M | 0.66M |
| E(3)-GNN | 11.32M | 11.32M |
| CrossAttentionFusion | 2.10M | 2.10M |
| ClassificationHead (Teacher) | 0.13M | 0.13M |
| **Teacher TOTAL** | **665.25M** | **92.92M** |

Table 3: Cross-attention fusion module

| Hyperparameter | Value |
|---|---|
| Attention heads | 8 |
| Embedding dimension | 512 |
| FFN intermediate dimension | 1024 |

Table 4: Classification head (shared by teacher and student)

| Hyperparameter | Value |
|---|---|
| Architecture | Linear(512→256)–ReLU–Linear(256→ $C$) |
| Dropout | 0.1 |

Table 5: Student model parameter counts.

| Module | Total params | Trainable params |
|---|---|---|
| ESM2-35M (backbone) | 33.38M | 33.38M |
| ESM2 projection | 0.26M | 0.26M |
| ClassificationHead (Student) | 0.13M | 0.13M |
| **Student TOTAL** | **33.77M** | **33.77M** |

#### S1.2 Training Procedure

**Phase 1 – Teacher training.** The teacher comprises ESM2-650M (`esm2_t33_650M_UR50D`) and an E(3)-equivariant graph neural network (EGNN) [3]. The EGNN satisfies full E(3) equivariance (rotations, translations, and reflections) by constructing edge messages solely from pairwise squared distances $\|x_i - x_j\|^2$ and node features, and updating $C_\alpha$ coordinates via radial-direction weighted aggregation. ESM2 parameters were frozen except for the final 4 transformer layers and the post-embedding layer normalisation, which were fine-tuned with a reduced learning rate. All models were optimised with AdamW [2]. Hyperparameters are listed in 6.

Table 6: Teacher training hyperparameters

| Hyperparameter | Value |
|---|---|
| Epochs | 10 |
| Batch size | 2 |
| Optimiser | AdamW |
| Learning rate (ESM2 unfrozen layers) | $1 \times 10^{-5}$ |
| Learning rate (GNN / Fusion / Head) | $1 \times 10^{-4}$ |
| Weight decay | $1 \times 10^{-2}$ |
| LR schedule | Cosine annealing ($T_{\max}$=10, $\eta_{\min}$=$10^{-6}$) |
| Gradient clipping | $\|\mathbf{g}\|_2 \leq 1.0$ |
| Model selection | Best validation F1 |

**Phase 2 – Knowledge distillation.** The student uses ESM2-35M (`esm2_t12_35M_UR50D`), using layer-12 representations of dimension 480 projected to 512. The distillation objective is

$$\mathcal{L} = \lambda_1 \mathcal{L}_{\text{logits}} + \lambda_2 \mathcal{L}_{\text{fusion}} + \mathcal{L}_{\text{task}}, \tag{1}$$

where $\mathcal{L}_{\text{task}}$ is cross-entropy, $\mathcal{L}_{\text{logits}} = \tau^2 \,\mathrm{KL}(\sigma(\mathbf{y}_t/\tau)\|\sigma(\mathbf{y}_s/\tau))$ is the temperature-scaled logits loss, and $\mathcal{L}_{\text{fusion}} = \mathrm{MSE}(\mathbf{z}_s, \mathbf{z}_t)$ aligns the student representation to the fused teacher embedding. We set $\lambda_1 = 1.0$, $\lambda_2 = 0.1$, and $\tau = 4.0$; the smaller $\lambda_2$ down-weights representation alignment relative to logits distillation. Remaining hyperparameters are listed in Table 7.

Table 7: Student distillation hyperparameters

| Hyperparameter | Value |
|---|---|
| Epochs | 10 |
| Batch size | 2 |
| Optimiser | AdamW |
| Learning rate | $1 \times 10^{-4}$ |
| Weight decay | $1 \times 10^{-2}$ |
| LR schedule | Cosine annealing ($T_{\max}$=10, $\eta_{\min}$=$10^{-6}$) |
| Gradient clipping | $\|\mathbf{g}\|_2 \leq 1.0$ |
| Distillation temperature $\tau$ | 4.0 |
| $\lambda_1$ (logits loss weight) | 1.0 |
| $\lambda_2$ (fusion loss weight) | 0.1 |
| Model selection | Best validation F1 |

#### S1.3 Data Partitioning and Reproducibility

GeoARG-DB was randomly partitioned (seed 42) into training (70%), validation (10%), and test (20%) subsets. Protein sequences longer than 1,022 residues were truncated at the sequence level; the corresponding PDB structures were truncated to the same length to preserve residue-level alignment between the sequence encoder and the graph neural network. The test set was evaluated only once after all hyperparameter decisions were finalised. All experiments were conducted on a single NVIDIA A100 GPU using PyTorch.

#### S1.4 GeoARG-DB Resistance Class Distribution

GeoARG-DB comprises 40,624 non-redundant ARG protein sequences spanning 36 resistance classes. Classes were harmonised from seven source databases using a rule-based normalisation pipeline; sequences that could not be assigned to a defined resistance class were retained as "unclassified". The complete per-class sequence counts are listed in Table 8.

Table 8: **Sequence counts per resistance class in GeoARG-DB.** Classes are sorted by sequence count (descending). Percentages are computed relative to the total of 40,624 sequences.

| **Rank** | **Resistance Class** | **Count** | **%** |
|---|---|---|---|
| 1 | beta-lactam | 10,482 | 25.8 |
| 2 | multidrug | 5,208 | 12.8 |
| 3 | cephalosporin | 4,380 | 10.8 |
| 4 | bacitracin | 4,267 | 10.5 |
| 5 | mls | 3,037 | 7.5 |
| 6 | carbapenem | 2,606 | 6.4 |
| 7 | aminoglycoside | 2,303 | 5.7 |
| 8 | unclassified | 1,877 | 4.6 |
| 9 | polymyxin | 1,380 | 3.4 |
| 10 | tetracycline | 1,093 | 2.7 |
| 11 | phenicol | 746 | 1.8 |
| 12 | fosfomycin | 736 | 1.8 |
| 13 | glycopeptide | 720 | 1.8 |
| 14 | quinolone | 663 | 1.6 |
| 15 | trimethoprim | 392 | 1.0 |
| 16 | sulfonamide | 272 | 0.7 |
| 17 | rifamycin | 117 | 0.3 |
| 18 | nitroimidazole | 44 | 0.1 |
| 19 | kasugamycin | 36 | 0.1 |
| 20 | fosmidomycin | 29 | 0.1 |
| 21 | aminocoumarin | 29 | 0.1 |
| 22 | spectinomycin | 28 | 0.1 |
| 23 | fusidic-acid | 27 | 0.1 |
| 24 | bleomycin | 25 | 0.1 |
| 25 | oxazolidinone | 24 | 0.1 |
| 26 | streptothricin | 21 | 0.1 |
| 27 | mupirocin | 18 | <0.1 |
| 28 | peptide | 17 | <0.1 |
| 29 | elfamycin | 13 | <0.1 |
| 30 | tetracenomycin-c | 8 | <0.1 |
| 31 | pleuromutilin | 8 | <0.1 |
| 32 | isoniazid | 5 | <0.1 |
| 33 | tiamulin | 4 | <0.1 |
| 34 | ethambutol | 4 | <0.1 |
| 35 | puromycin | 3 | <0.1 |
| 36 | hygromycin | 2 | <0.1 |

| | | |
|---|---|---|
| **Total** | **40,624** | **100.0** |

MLS: macrolide–lincosamide–streptogramin. "Unclassified" includes sequences from source databases lacking a defined resistance-class annotation, sequences with conflicting cross-database annotations, and entries for which the rule-based normalisation pipeline could not assign a canonical class. The bacitracin category (4,267 sequences, 10.5%) is dominated by HMD-ARG-derived entries (3,987 of 4,267; 93.4%), which annotates undecaprenyl pyrophosphate phosphatase (UppP) family proteins as bacitracin-resistance determinants. This annotation reflects HMD-ARG's broader inclusion of target-protection and membrane-remodelling proteins; other databases apply a narrower criterion and contribute only 280 sequences to this class. This source-specific enrichment should be considered when interpreting model performance on the bacitracin class.

#### S1.5 ResFinderFG Identity Bin Statistics

Table 9 summarises the number of ResFinderFG v2.0 sequences in each sequence identity bin relative to GeoARG-DB-Train.

Table 9: Number of ResFinderFG v2.0 sequences per identity bin.

| Identity to GeoARG-DB (%) | Sequences |
|---|---|
| 0–40 | 1,083 |
| 40–60 | 1,233 |
| 60–100 | 1,147 |
| **Total** | **3,463** |

#### S1.6 MCR-like Dataset Construction Statistics

Table 10: Sequence counts at each stage of *mcr*-like dataset construction.

| Source / Processing step | Sequences |
|---|---|
| tBLASTn hits (*mcr*-1, `WP_163397051.1`, top 5,000) | 14,197[a] |
| tBLASTn hits (*mcr*-4, `WP_099156046.1`, top 5,000) | 19,401[a] |
| MicroBIGG-E *mcr*-labelled entries | 17,071 |
| EptA/B/C (*mcr*-like) sequences | 324 |
| Total after merging and deduplication | 29,352 |
| After length filtering (100–900 aa) for phylogeny | 29,256 |
| *Canonical mcr* (source metadata) | 17,071 |
| *Expanded mcr* (MRCA-based) | 19,163 |
| Non-*mcr* background | 10,093 |

[a] CDS features extracted from GenBank records overlapping the tBLASTn-aligned region; one hit may yield multiple CDS entries.

#### S1.7 MCR-like Phylogenetic Tree Construction

Multiple sequence alignment was performed using MAFFT (FFT-NS-i iterative refinement mode) [5]. Prior to tree inference, alignment columns with site occupancy below 0.20 were removed, and sequences retaining more than 95% gap characters after column trimming were discarded, yielding 29,256 sequences for phylogenetic reconstruction.

A maximum-likelihood phylogenetic tree was inferred using FastTree (v2.2.0) [6] under the LG substitution model (Le-Gascuel 2008) with the CAT approximation (20 rate categories). Branch support was estimated using the SH-like test with 1,000 replicates. Tree search employed the default NNI and SPR (2 rounds, range 10) heuristics:

```
FastTree -lg <trimmed.mafft.fasta> > <output.fasttree.nwk>
```

To define the expanded *mcr* set, the most recent common ancestor (MRCA) of each *mcr* subtype (*mcr*-1 through *mcr*-10) was identified in the inferred tree using the 17,071 MicroBIGG-E source-labelled sequences as anchors. All descendant leaves of each subtype MRCA were incorporated into the expanded *mcr* set, yielding 19,163 expanded *mcr* sequences and 10,093 non-*mcr* background sequences (Table 10).

##### S1.8 GMGC dataset and candidate

Table 11: GMGC gut sequence filtering statistics.

| **Filtering step** | **Sequences** |
|---|---|
| GMGC gut catalog (total ORFs) | 4,644,769 |
| After <25% identity filter | 111,202 |
| After retaining unannotated only | 49,707 |
| GeoARG high-confidence ($P > 0.8$) | 1,485 |

#### S2. Extended Benchmark Results

##### S2.1 UniProt Benchmark

Table 12: Binary ARG identification on the UniProt benchmark (GeoARG-DB-Test positives + UniProt-derived negatives).

| **Model** | **Accuracy** | **Sensitivity** | **Specificity** | **MCC** | **AUROC** |
|---|---|---|---|---|---|
| DeepARG | 0.8697 | 0.7410 | 0.9960 | 0.7637 | 0.8692 |
| ARGNet | 0.9406 | 0.9385 | 0.9422 | 0.8793 | 0.9674 |
| GeoARG | **0.9992** | **0.9991** | **0.9993** | **0.9984** | **0.9999** |

##### S2.2 ORF Benchmark

Table 13: Binary ARG identification on the ORF benchmark across four input settings. LSnt/SSnt: long/short nucleotide sequences; LSaa/SSaa: long/short amino acid sequences. Best results per setting are in **bold**.

| **Setting** | **Model** | **Accuracy** | **Sensitivity** | **Specificity** | **MCC** | **AUROC** |
|---|---|---|---|---|---|---|
| LSnt | DeepARG | 0.9200 | 0.8718 | 0.9527 | 0.8253 | 0.9176 |
| | ARGNet | 0.8343 | 0.8553 | 0.8119 | 0.6683 | 0.9075 |
| | GeoARG | **0.9931** | **0.9881** | **0.9983** | **0.9862** | **0.9989** |
| SSnt | DeepARG | 0.9236 | 0.8556 | 0.9945 | 0.8563 | 0.9263 |
| | ARGNet | 0.9195 | 0.8614 | 0.9799 | 0.8455 | 0.9282 |
| | GeoARG | **0.9472** | **0.9131** | **0.9827** | **0.8969** | **0.9826** |
| LSaa | DeepARG | 0.8390 | 0.7102 | 0.9724 | 0.7050 | 0.8439 |
| | ARGNet | 0.9315 | 0.8789 | 0.9151 | 0.8634 | 0.9706 |
| | GeoARG | **0.9987** | **0.9979** | **0.9994** | **0.9973** | **0.9998** |
| SSaa | DeepARG | 0.8096 | 0.6211 | 0.9991 | 0.6695 | 0.8102 |
| | ARGNet | 0.9328 | 0.8789 | 0.9870 | 0.8708 | 0.9373 |
| | GeoARG | **0.9443** | **0.9220** | **0.9770** | **0.8895** | **0.9816** |

#### S3. GeoARG-DB Composition

Table 14: Number of sequences contributed by each source database. Raw counts are after merging; the final total reflects removal of SNP-mediated resistance entries and CD-HIT deduplication at 100% identity.

| **Database** | **Raw sequences** |
|---|---|
| CARD | 6,627 |
| ResFinder | 3,218 |
| AMRFinder+ | 6,081 |
| MEGARes | 9,044 |
| DeepARG-DB | 14,933 |
| HMD-ARG-DB | 8,285 |
| ARGNet-DB | 10,583 |
| Total (merged) | 58,771 |
| After SNP exclusion | 55,340 |
| After deduplication | 40,624 |

#### S4. ESMfold and AlphaFold3 Confidence Scores

Protein structures were predicted using ESMFold [1]. Per-residue pLDDT scores were extracted from the B-factor field of each predicted PDB file. The distributions are highly consistent between the training and test sets, confirming that the random partition did not introduce systematic bias in structural confidence. Residues with pLDDT $< 70$ were excluded during geometric feature construction, accounting for 13.71% and 13.34% of residues in the training and test sets, respectively.

Table 15: Per-residue pLDDT score statistics for ESMFold-predicted structures in the training and test sets.

| **Statistic** | **Train** | **Test** |
|---|---|---|
| Mean pLDDT | 85.51 | 85.76 |
| Median pLDDT | 94.90 | 94.93 |
| Standard deviation | 22.01 | 21.71 |
| Residues with pLDDT $< 70$ (%) | 13.71 | 13.34 |
| Residues with pLDDT $\geq 90$ (%) | 68.91 | 69.12 |

AlphaFold3 [4] was used to predict protein–ligand complex structures by co-folding each candidate protein with ampicillin. Confidence metrics for the representative beta-lactamase candidate GMGC10.207_616_678.UNKNOWN are summarised below. An ipTM $\geq 0.8$ indicates high-confidence interface prediction; the absence of steric clashes and disordered regions further supports the reliability of the predicted binding pose. The low per-chain pTM for the ligand (0.29) is expected for small molecules, which lack the secondary structure that pTM is calibrated for; the interface-level ipTM (0.86) is the appropriate confidence metric for protein–ligand complexes.

Table 16: AlphaFold3 confidence scores for the predicted protein–ampicillin complex of candidate GMGC10.207_616_678.UNKNOWN.

| Metric | Description | Value |
|---|---|---|
| ipTM | Interface confidence (protein–ligand) | 0.86 |
| pTM | Overall structural confidence | 0.95 |
| Ranking score | AF3 composite ranking score | 0.88 |
| Chain pTM (protein) | Per-chain pTM, protein | 0.96 |
| Chain pTM (ligand) | Per-chain pTM, ligand | 0.29[a] |
| Chain ipTM (protein) | Per-chain ipTM, protein | 0.86 |
| Chain ipTM (ligand) | Per-chain ipTM, ligand | 0.86 |
| Fraction disordered | Proportion of disordered residues | 0.00 |
| Steric clashes | Predicted atomic clashes | None |

[a] The low per-chain pTM for the ligand (0.29) is expected for small molecules, which lack the secondary structure that pTM is calibrated for. The interface-level ipTM (0.86) is the appropriate confidence metric for protein–ligand complexes.

### S5. Pfam Domain Enrichment Analysis of High-Confidence ARG Candidates

Pfam domain enrichment was assessed by comparing 1,485 high-confidence ARG candidates ($P > 0.8$) against 3,000 low-confidence background sequences ($P < 0.2$) drawn from the same unannotated GMGC gut sequence space. Fisher's exact test was applied to $2 \times 2$ contingency tables, odds ratios were computed with a pseudocount of 0.5, and multiple testing correction used the Benjamini–Hochberg procedure. Of 148 Pfam features tested, 36 were significantly enriched and 66 were significantly depleted (FDR $< 0.05$); the remainder were non-significant. The observed count of 105 significant features (enriched + depleted) far exceeds the random expectation estimated by 50 label-permutation replicates (mean = 0.08 significant features per replicate; maximum observed = 2). All 36 enriched features are listed in Table 17.

Table 17: **Pfam features significantly enriched in high-confidence ARG candidates relative to low-confidence background sequences.** Features are sorted by FDR (ascending). Odds ratios (OR) are computed with a pseudocount of 0.5; $\log_2$OR $> 0$ indicates enrichment in the candidate set. Features with zero background counts are marked with †.

| Pfam Feature | Resistance / Functional Relevance | $\log_2$ OR | FDR |
|---|---|---|---|
| *High-confidence enrichment (FDR $< 10^{-8}$)* | | | |
| HisKA | Histidine kinase A domain (VanS-type sensors) | 2.96 | $2.7 \times 10^{-133}$ |
| HATPase_c | Histidine kinase ATPase domain | 2.75 | $1.7 \times 10^{-92}$ |
| Peptidase_M56† | Membrane-anchored metallopeptidase | 9.72 | $7.1 \times 10^{-90}$ |
| HAMP | HAMP linker domain (signal transduction) | 3.54 | $1.0 \times 10^{-63}$ |
| CAT† | Chloramphenicol acetyltransferase | 6.99 | $1.1 \times 10^{-40}$ |
| SpaA | Lantibiotic-resistance associated | 6.01 | $5.5 \times 10^{-33}$ |
| SdrD_B† | Surface-anchored MSCRAMM domain | 5.10 | $1.9 \times 10^{-16}$ |
| Acyl_transf_3† | Acyltransferase family 3 | 7.06 | $9.3 \times 10^{-15}$ |
| Transpeptidase | DD-transpeptidase / PBP active site | 5.56 | $2.1 \times 10^{-14}$ |
| Beta-lactamase2 | Class B metallo-beta-lactamase | 4.78 | $6.8 \times 10^{-13}$ |
| HH_EMRA | EmrA-family efflux membrane fusion protein | 5.38 | $1.3 \times 10^{-12}$ |
| *Moderate enrichment (FDR $10^{-8}$–$10^{-3}$)* | | | |

Table 17 (continued)

| **Pfam Feature** | **Resistance / Functional Relevance** | $\log_2$ **OR** | **FDR** |
|---|---|---|---|
| HH_YBHG† | YbhG homologue, lipopolysaccharide transport | 6.52 | $4.2 \times 10^{-10}$ |
| APH | Aminoglycoside phosphotransferase | 3.45 | $6.7 \times 10^{-9}$ |
| YadA_stalk† | Adhesin/outer-membrane stalk domain | 6.31 | $1.0 \times 10^{-8}$ |
| bMG3† | Beta-barrel membrane protein family | 6.06 | $2.6 \times 10^{-7}$ |
| VanZ† | VanZ glycopeptide resistance protein | 5.54 | $5.5 \times 10^{-5}$ |
| Anti-Pycsar_Apyc1† | Anti-Pycsar defence system | 5.54 | $5.5 \times 10^{-5}$ |
| Lactamase_B | Metallo-beta-lactamase fold | 2.85 | $5.5 \times 10^{-5}$ |
| NTP_transf_2† | Nucleotidyl transferase (aminoglycoside) | 5.26 | $4.3 \times 10^{-4}$ |
| PP-binding† | Phosphopantetheine-binding (NRPS/PKS) | 5.26 | $4.3 \times 10^{-4}$ |
| *Nominal enrichment (FDR 0.001–0.05)* | | | |
| AadA_C† | Aminoglycoside 3'-adenyltransferase C-term | 4.92 | $3.6 \times 10^{-3}$ |
| Peptidase_S11† | DD-peptidase / PBP family | 4.92 | $3.6 \times 10^{-3}$ |
| HATPase_c_2 | Histidine kinase-like ATPase variant | 3.08 | $4.0 \times 10^{-3}$ |
| NOMO1-like_2nd | Signal-peptide processing related | 1.53 | $6.1 \times 10^{-3}$ |
| Acetyltransf_1 | Acetyltransferase (aminoglycoside/phenicol) | 1.44 | $9.2 \times 10^{-3}$ |
| Choline_kinase† | Choline/ethanolamine kinase fold | 4.71 | $9.2 \times 10^{-3}$ |
| HATPase_c_3 | Histidine kinase-like ATPase variant 3 | 2.37 | $1.1 \times 10^{-2}$ |
| DUF5780† | Domain of unknown function | 4.47 | $2.3 \times 10^{-2}$ |
| Polbeta† | DNA polymerase beta-like nucleotidyltransferase | 4.47 | $2.3 \times 10^{-2}$ |
| TetR_C_39† | TetR-family C-terminal domain variant | 4.47 | $2.3 \times 10^{-2}$ |
| Adenyl_transf† | Adenylyltransferase (aminoglycoside) | 4.47 | $2.3 \times 10^{-2}$ |
| TetR_N | TetR N-terminal helix-turn-helix domain | 1.07 | $2.4 \times 10^{-2}$ |
| Acetyltransf_7 | Acetyltransferase variant 7 | 1.26 | $4.4 \times 10^{-2}$ |
| Acetyltransf_3 | Acetyltransferase variant 3 | 2.29 | $4.4 \times 10^{-2}$ |
| AMP-binding_C_3 | AMP-binding protein C-terminal domain 3 | 2.29 | $4.4 \times 10^{-2}$ |

† Feature absent in background sequences (count = 0); odds ratio computed with pseudocount 0.5. OR: odds ratio; FDR: false discovery rate (Benjamini–Hochberg correction).

### S6. High-Confidence Candidates Are Evolutionarily Divergent yet Functionally Conserved

This section provides methodological details for the analyses presented in the main-text subsection "High-confidence candidates are evolutionarily divergent yet functionally conserved" and in Figs. 5–6.

#### S6.1 Candidate Filtering and Class Distribution

From the 49,707 unannotated GMGC gut sequences, 1,485 were classified as high-confidence ARG candidates (prediction probability $P > 0.8$). Analysis was restricted to the six resistance classes for which interpretable functional annotations (Pfam domains or catalytic motifs) were available. Class labels were normalised to lowercase; "beta-lactamase" predictions were mapped to "beta-lactam" for consistency with GeoARG-DB nomenclature. The per-class candidate counts are listed in Table 18.

Table 18: High-confidence ARG candidates ($P > 0.8$) per resistance class.

| **Resistance class** | **Candidates** |
|---|---|
| Glycopeptide | 605 |
| Beta-lactam | 590 |
| MLS | 98 |
| Phenicol | 90 |
| Aminoglycoside | 67 |
| Tetracycline | 35 |
| **Total** | **1,485** |

#### S6.2 Taxonomic Assignment of ARG Candidates

**Phylum-level taxonomy from GMGC metadata (Fig. 5a).** Phylum-level taxonomic assignments were derived from the GMGC v10 pre-computed taxonomy table (`metadata_GMGC10.taxonomy.tsv`), which provides NCBI Taxonomy assignments at multiple ranks for each unigene. When multiple ranks were available for a single sequence, the most specific rank was retained using the following priority order: species > genus > family > order > class > phylum > superkingdom. Sequences absent from the taxonomy table or lacking any assigned rank were labelled "Unassigned." For visualisation, the four most abundant phyla (by total candidate count across all resistance classes) were displayed individually; all remaining phyla were merged into an "Other" category.

**Genus-level taxonomy from DIAMOND (Fig. 5b).** Genus-level best-hit annotation was obtained by searching the 1,485 candidate protein sequences against a bacterial subset of UniProt (UniProtKB, downloaded 10 November 2025; taxonomy ID 2, excluding entries with computational function annotation) using DIAMOND blastp v2.1 with the `--sensitive` flag:

```
diamond blastp \
  --query candidates.faa \
  --db uniprot_bacteria.dmnd \
  --out top1.tsv \
  --outfmt 6 qseqid sseqid pident length mismatch \
             gapopen qstart qend sstart send evalue bitscore \
  --max-target-seqs 1 \
  --sensitive \
  --threads 8
```

The default DIAMOND e-value threshold ($10^{-1}$) was applied. For each candidate with at least one hit, the genus was extracted as the first whitespace-delimited token of the UniProt "Organism" field after removing parenthetical strain designations. Candidates with no DIAMOND hit were labelled "No hit." The four most abundant genera (excluding "No hit") were displayed individually; all others were merged into "Other."

**Environment assignment.** Environment-of-origin annotations were obtained from the GMGC v10 pre-computed metadata table (`metadata_GMGC10.unigene-environment.tsv`). Each unigene carries a ternary score (0, 1, or 2) across multiple habitat categories (e.g., gut, soil, freshwater). Presence was defined as a score $\geq 1$; strong association was defined as a score of exactly 2. When a sequence was associated with multiple environments at equal maximum strength, it was labelled "multi." Sequences absent from the environment table were labelled "unassigned."

#### S6.3 Pfam Domain Annotation and Support Classification

Pfam domain annotation was performed using HMMER v3.3 hmmscan against the Pfam-A database (release 35) with an independent e-value (i-Evalue) threshold of $1 \times 10^{-5}$ per domain hit.

A candidate sequence was classified as *Pfam-supported* if it carried at least one Pfam domain from a manually curated, class-specific whitelist of canonical resistance-associated domains (Table 19). The whitelist was constructed by selecting Pfam families with established mechanistic links to each resistance class. Sequences with Pfam hits that did not match the whitelist for their predicted resistance class were classified as *not supported*.

Table 19: Class-specific Pfam domain whitelists used to assess functional support of ARG candidates.

| **Resistance class** | **Accepted Pfam domains** |
|---|---|
| Beta-lactam | Beta-lactamase, Beta-lactamase2, Lactamase_B |
| Phenicol | CAT |
| Aminoglycoside | APH, Adenyl_transf, AadA_C, Acetyltransf_1, Acetyltransf_3, Acetyltransf_8, NTP_transf_2, Antibiotic_NAT |
| Glycopeptide | HATPase_c, HisKA, HAMP, Response_reg, VanZ |
| Tetracycline | TetR_N, TetR_C_39 |
| MLS | ABC_tran, YknX_C, BSH_RND, HH_EMRA, Beta-barrel_YknX, CzcB_C, BSH_CzcB, HH_YBHG |

#### S6.4 Beta-Lactamase Catalytic Motif Detection

The three canonical serine beta-lactamase catalytic motifs were detected using regular-expression searches on the full-length amino acid sequences of beta-lactam candidates:

- **SXXK**: `S[A-Z]{2}K`
- **SXN**: `S[A-Z]N`
- **KTG**: `KTG`

where `[A-Z]` denotes any standard amino acid. No positional constraints were applied; a motif was scored as present if the pattern occurred at least once anywhere in the sequence. A candidate was classified as retaining catalytic architecture if it contained at least two of the three motifs ($\geq 2$ motifs).

#### S6.5 Phylogenetic Tree Construction

Both the phenicol (Fig. 5d) and beta-lactamase (Fig. 4c) phylogenetic trees were constructed using the same pipeline. In each case, known ARG sequences from GeoARG-DB were manually curated to include representative members of the target class, pooled with the corresponding high-confidence candidates ($P > 0.8$), and clustered at 50% sequence identity using CD-HIT v4.8.1 (`cd-hit -c 0.5 -n 2`).

Multiple sequence alignment was performed using MAFFT (FFT-NS-i iterative refinement) [5]. A maximum-likelihood tree was inferred using FastTree v2.2.0 [6] under the LG substitution model with the CAT rate approximation (20 categories). The resulting trees were visualised using iTOL v6.

**Phenicol tree (Fig. 5d).** Known phenicol ARG sequences from GeoARG-DB were pooled with the 20 high-confidence phenicol candidates. After clustering at 50% identity, 78 representative sequences were retained (58 known, 20 candidates). Inner-ring annotations indicate sequence type (known ARG vs. predicted candidate); outer-ring annotations indicate CAT domain presence as determined by HMMER hmmscan (i-Evalue $< 10^{-5}$).

**Beta-lactamase tree (Fig. 4c).** Representative class A beta-lactamase sequences were manually selected from GeoARG-DB to capture the major known subfamilies. These were pooled with the high-confidence beta-lactam candidates, clustered at 50% identity using CD-HIT, and processed through the MAFFT–FastTree pipeline described above.

##### S6.6 Foldseek Structural Alignment

Structural alignments were performed using Foldseek v9 with the `easy-search` workflow against PDB-format structures predicted by ESMFold. Default alignment parameters were used.

For the representative beta-lactamase candidate (GMGC10.207_616_678.UNKNOWN), the reference structure was the ESMFold-predicted model of a class A beta-lactamase (`WP_064935627.1`). Motif-level C$\alpha$ RMSD was computed over residues corresponding to the SXXK, SXN, and KTG motifs after structural superposition by TM-align.

For the glycopeptide candidate (`GMGC10_052_200_458.UNKNOWN`), the reference was VanS-D (`WP_063856738.1`), with pairwise C$\alpha$ distances computed over the conserved H-box, N-box, and G-box motifs.

##### S6.7 Counterfactual Motif Perturbation Analysis

To interpret which input features drive GeoARG predictions, we performed *in silico* counterfactual perturbation analysis on 20 high-confidence beta-lactam candidates selected on the basis of prediction confidence ($P > 0.80$), structural prediction quality (pLDDT $> 90$), and complete retention of all three canonical catalytic motifs (SXXK, SXN, and KTG).

Canonical catalytic-motif residues were individually or pairwise replaced with alanine. The following perturbation conditions were evaluated:

1. Single-motif disruption: SXXK only; SXN only; KTG only.
2. Pairwise disruption: SXXK + SXN; SXXK + KTG; SXN + KTG.

Each condition was applied independently to all 20 sequences, yielding $n = 20$ observations per group. For each perturbed sequence, the modified amino acid sequence was passed directly through the GeoARG student inference pipeline without regenerating protein structures, as the student model operates on sequence input only.

#### S7. Molecular Dynamics Simulation Details

MD simulations were performed using GROMACS with the CHARMM36m force field and TIP3P explicit water model. The simulation protocol is summarised in Table 20.

Table 20: Molecular dynamics simulation parameters.

| **Parameter** | **Value** |
|---|---|
| Force field | CHARMM36m |
| Water model | TIP3P |
| Production time | 200 ns |
| Time step | 2 fs |
| Temperature | 310 K (V-rescale) |
| Pressure | 1 bar (C-rescale) |
| Electrostatics | PME |
| VdW cutoff | 1.2 nm |
| Energy minimisation | Steepest descent |
| NVT equilibration | 5 ns |
| Analysis tools | GROMACS, PyMOL |
